# Reproducible single cell annotation of programs underlying T-cell subsets, activation states, and functions

**DOI:** 10.1101/2024.05.03.592310

**Authors:** Dylan Kotliar, Michelle Curtis, Ryan Agnew, Kathryn Weinand, Aparna Nathan, Yuriy Baglaenko, Yu Zhao, Pardis C. Sabeti, Deepak A. Rao, Soumya Raychaudhuri

**Author notes:** These authors contributed equally (Dylan Kotliar, Michelle Curtis). Address correspondence to: Soumya Raychaudhuri, 77 Avenue Louis Pasteur, Harvard New Research Building, Suite 250D Boston, MA 02446, USA.

## Abstract

T-cells recognize antigens and induce specialized gene expression programs (GEPs) enabling functions including proliferation, cytotoxicity, and cytokine production. Traditionally, different classes of helper T-cells express mutually exclusive responses – for example, Th1, Th2, and Th17 programs. However, new single-cell RNA sequencing (scRNA-Seq) experiments have revealed a continuum of T-cell states without discrete clusters corresponding to these subsets, implying the need for new analytical frameworks. Here, we advance the characterization of T-cells with T-CellAnnoTator (TCAT), a pipeline that simultaneously quantifies pre-defined GEPs capturing activation states and cellular subsets. From 1,700,000 T-cells from 700 individuals across 38 tissues and five diverse disease contexts, we discover 46 reproducible GEPs reflecting the known core functions of T-cells including proliferation, cytotoxicity, exhaustion, and T helper effector states. We experimentally characterize several novel activation programs and apply TCAT to describe T-cell activation and exhaustion in Covid-19 and cancer, providing insight into T-cell function in these diseases.

## Introduction

Canonically, T-cells are classified by membership in a hierarchy of discrete, mutually exclusive subsets associated with key transcription factors and surface markers. For example, expression of γδ or αꞵ T-cell receptors and CD4 or CD8 co-receptors divide T-cells into subsets recognizing different major histocompatibility complex (MHC) molecules. CD45 isoform and L-selectin expression subdivides naive and memory subsets. CD4 memory cells are further subcategorized into helper subsets, including Th1, Th2, and Th17, with distinct cytokine profiles upon activation^1^.

Emerging evidence conflicts with this canonical model. T-cell states vary continuously^2^, combine additively within a cell^3^, and have plasticity in response to stimuli^4^. This may explain why single-cell RNA sequencing (scRNA-Seq) typically shows a continuum of T-cell states without well-delineated clusters corresponding to discrete subsets^5,6^. Even with incorporation of pre-defined surface protein markers based on cellular indexing of transcriptomes and epitopes by sequencing (CITE-seq)^7^, unbiased clustering does not yield canonical discrete T-helper subsets^8^. Rather, scRNA-Seq has highlighted untraditional cell populations including cytotoxic CD4+ cells^9^, CD8+ regulatory T-cells^10^ and Th1/Th17 cells^11^, consistent with the growing recognition of non discrete T-cell states.

While hard clustering is the predominant scRNA-Seq analysis technique, it has key limitations when cell states are not discrete or mutually exclusive. A cell’s transcriptome reflects its complex identity through expression of multiple gene expression programs (GEPs) that reflect lineage, activation states, and lifecycle processes^12^. However, hard clustering forces cells into discrete groups that cannot easily reflect the multiplicity of GEPs they express. For example, proliferating cells from multiple subsets may cluster together, obscuring information about their subset. Hard clustering also cannot directly model continuous expression trajectories and instead arbitrarily discretizes cells into distinct clusters.

Component-based models like non-negative matrix factorization (NMF), hierarchical Poisson factorization, and SPECTRA can overcome some of these limitations of hard clustering^5,13–16^. These methods model GEPs as vectors of expression values for each gene, and cells as weighted mixtures of GEPs. Unlike Principal Component Analysis (PCA), NMF components have been shown to correspond to biologically distinct GEPs^14^. Thus, NMF can capture instances where multiple GEPs reflecting cell-type and other functional states additively contribute to a cell’s transcriptome. Furthermore, unlike cluster assignments, GEP vectors may be able to serve as a fixed coordinate system onto which new datasets can be projected, enabling reproducible comparison of GEP activity across biological contexts. Previous analyses of T-cells using component-based models have already recognized GEPs associated with T-cell activation^5^ and exhaustion^15^.

We argue that scaling these approaches may further elucidate T-cell biology. First, most previous analyses have only analyzed T-cells from a small number of individual donors in a limited set of biological contexts. As a result, they have identified a modest number of GEPs. Moreover, it is essential to demonstrate the possibility of transferring GEPs identified in one dataset to new datasets. For example, it remains unclear whether reference GEPs learned in one dataset can accurately infer cell subsets, T-cell receptor (TCR)-dependent activation, and proliferation status for cells in a new dataset.

Here, we present CellAnnoTator (*CAT, pronounced starCAT), an approach to score cells based on a fixed, multidataset catalog of GEPs from any tissues or cell-type (indicated by the wildcard character “*”). We develop a catalog of GEPs reflecting the breadth of subsets, activation states, and functions within T-cells by applying consensus NMF (cNMF)^14^, a validated implementation of NMF, to 7 scRNA-Seq datasets, spanning 1.7 million T-cells across 38 human tissues^6,8,11,17–20^. We observe striking concordance of many GEPs across contexts. After combining analogous GEPs, we define a final catalog of 46 consensus GEPs (cGEPs) capturing diverse features of T-cells (**Figure 1A**). We demonstrate *CAT by accurately inferring T-cell subsets in query datasets and quantifying rates of TCR-dependent activation and exhaustion in Covid-19 and cancer.

**Figure 1.**
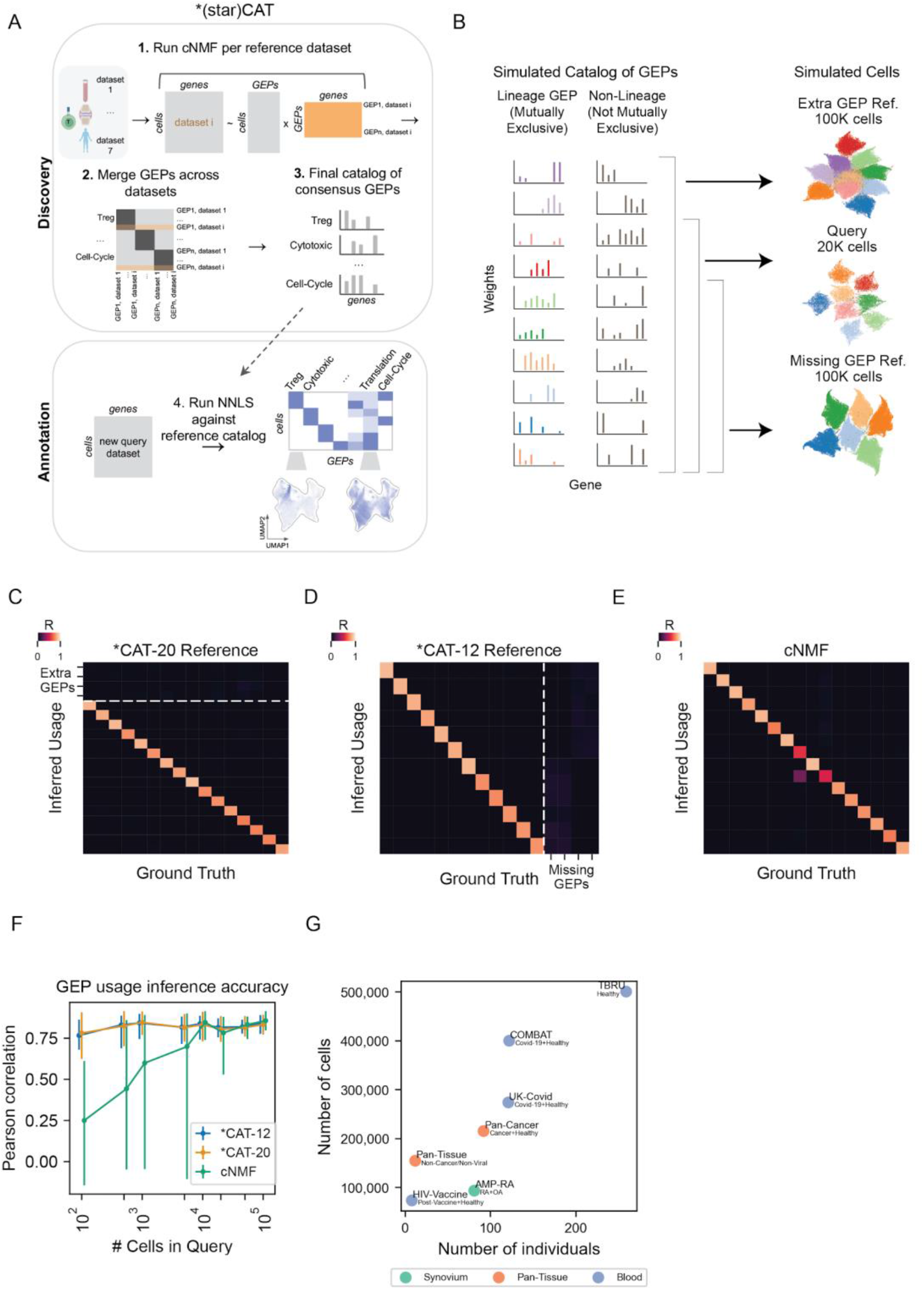
Overview of CellAnnoTator (*CAT). (A) Schematic of the *CellAnnoTator (*CAT) pipeline. (B) Schematic of simulation strategy (left) with resulting Uniform Manifold Approximation and Projection (UMAP) plot (right). Cells are colored by lineage gene expression program (GEP). (C-E) Pearson correlation of ground truth simulated usages of each GEP (columns) vs inferred usages (rows) for *CAT with the 20 GEP reference (C), *CAT with the 12 GEP reference (D) or cNMF of the query with 16 inferred components (E). (F) Pearson correlation of ground truth and inferred usages by *CAT and cNMF for different query dataset sizes. Marker represents mean and error bars represent range. (G) Summary of reference datasets including number of individual donors (x-axis), number of cells (y-axis), and tissue source (dot color). Phenotypes are listed below the dataset names.

## Results

### 1. Annotating cells with pre-defined gene expression programs

We first augmented the published cNMF algorithm to enhance GEP discovery, which is the first step of *CAT (**Figure 1A –top**). cNMF mitigates the randomness of individual NMF runs by repeating NMF with multiple seeds and combining the results into robust estimates^14^. It outputs GEP spectra, with gene weights for each GEP, and usages, reflecting the GEP’s weighted contribution to each cell. For our approach, it was essential to amalgamate the inferred GEP spectra from multiple datasets. However, we found that dataset-specific batch effects could hinder the identification of reproducible GEPs. Most batch correction methods are not compatible with cNMF since they create many negative values or correct low-dimensional embeddings rather than gene-level data. We therefore used Harmony^21^, with modifications to produce non-negative values for gene-level data rather than principal components. We also adapted cNMF to incorporate surface proteins into the final spectra to aid in GEP interpretation without impacting GEP discovery (**Methods**).

Next, we developed *CAT to enable GEPs learned in a reference dataset to be transferred to previously unseen “query” datasets. Whereas cNMF simultaneously learns GEPs and scores their usage in each cell’s transcriptional profile, *CAT addresses the independent problem of quantifying the usages of a fixed set of GEPs in a new dataset, using non-negative least squares (NNLS) regression, similar to NMFproject^13^. The result is a vector of usages for each cell representing the relative contribution of each GEP to the cell’s profile (**Figure 1A – bottom**).

Using NNLS to refit GEPs as we do with *CAT provides significant advantages over direct applications of cNMF or other matrix factorizations. First, *CAT uses a fixed set of GEPs from a reference, instead of discovering GEPs *de novo* in the query. Thus, it provides a consistent representation of cell states that can be compared across different datasets and biological contexts. Second, *de novo* cNMF might miss GEPs that are active in small numbers of cells, whereas *CAT can characterize activity in a query dataset with relatively few cells. Finally, *CAT is significantly faster to run than cNMF.

We conducted simulations to benchmark *CAT in scenarios where the reference and query datasets have only partially overlapping GEPs (**Methods**). We simulated two reference datasets of 100,000 cells and a query dataset of 20,000 cells. Each cell could express up to eleven GEPs, including one of ten mutually exclusive subset GEPs and up to ten non-subset GEPs. One reference dataset included all 16 GEPs in the query data as well as four additional GEPs. The other reference dataset was missing four GEPs present in the query (**Figure 1B**). We then learned GEPs from each reference dataset with cNMF and fit them to the query using *CAT. The reference and query datasets shared only 90% of genes in common, as datasets rarely share all genes.

*CAT accurately inferred the usage of GEPs that overlapped between the reference and query datasets (Pearson R>0.7) (**Figure 1C-D)**. *CAT had low predicted usage of the extra GEPs in the reference panel that were not in the query dataset (**Figure S1A**). Surprisingly, *CAT obtained better concordance with the simulated ground truth GEP usages than direct application of cNMF to the query (**Figure 1E)**. This is striking because the reference GEPs had extra or missing GEPs relative to the query, and were learned on different datasets, so could incorporate dataset-specific noise. We hypothesized that *CAT’s increased performance reflected the larger reference datasets enabling more accurate GEP inference. We confirmed this by simulating multiple query datasets with between 100 and 100,000 cells. While cNMFs performance declined for small query datasets, *CATs remained constant, demonstrating that *CAT can out-perform cNMF when the reference is larger than the query (**Figure 1F**).

### 2. Gene expression programs for T-cell annotation

We next developed a catalog of GEPs to capture T-cell states; combining these GEPs with the *CAT algorithm yields T-CellAnnoTator (TCAT). We analyzed T-cells from 7 diverse datasets including blood and tissues from healthy individuals or individuals with Covid-19, cancer, rheumatoid arthritis, or osteoarthritis (**Figure 1G).** After stringent quality control, there were 1.7 million cells from 905 samples from 695 individuals in our analysis. To preserve dataset-specific GEPs, we applied cNMF to each batch-corrected dataset independently (**Supplementary item 1, Methods**).

We observed that GEPs were reproducible across the datasets. To quantify this, we clustered highly correlated GEPs found in different datasets (**Methods**). Assuming that correlated dataset-specific GEPs represented the same biological state, we defined a consensus gene expression program (cGEP) as the average of a GEP cluster. Nine cGEPs derived from a cluster of GEPs from all seven datasets (Average Pearson R=0.81) and 49 cGEPs derived from a cluster of GEPs from two or more datasets (Average Pearson R=0.74) (**Figure 2A-B, S1B**). Between 68.4% and 96.8% of GEPs identified in each of the seven reference datasets clustered with at least one GEP from another reference, suggesting high reproducibility. By contrast, gene expression principal components showed limited concordance between pairs of datasets, suggesting they reflect more dataset-specific signals^14^ (**Figure S1C**).

**Figure 2.**
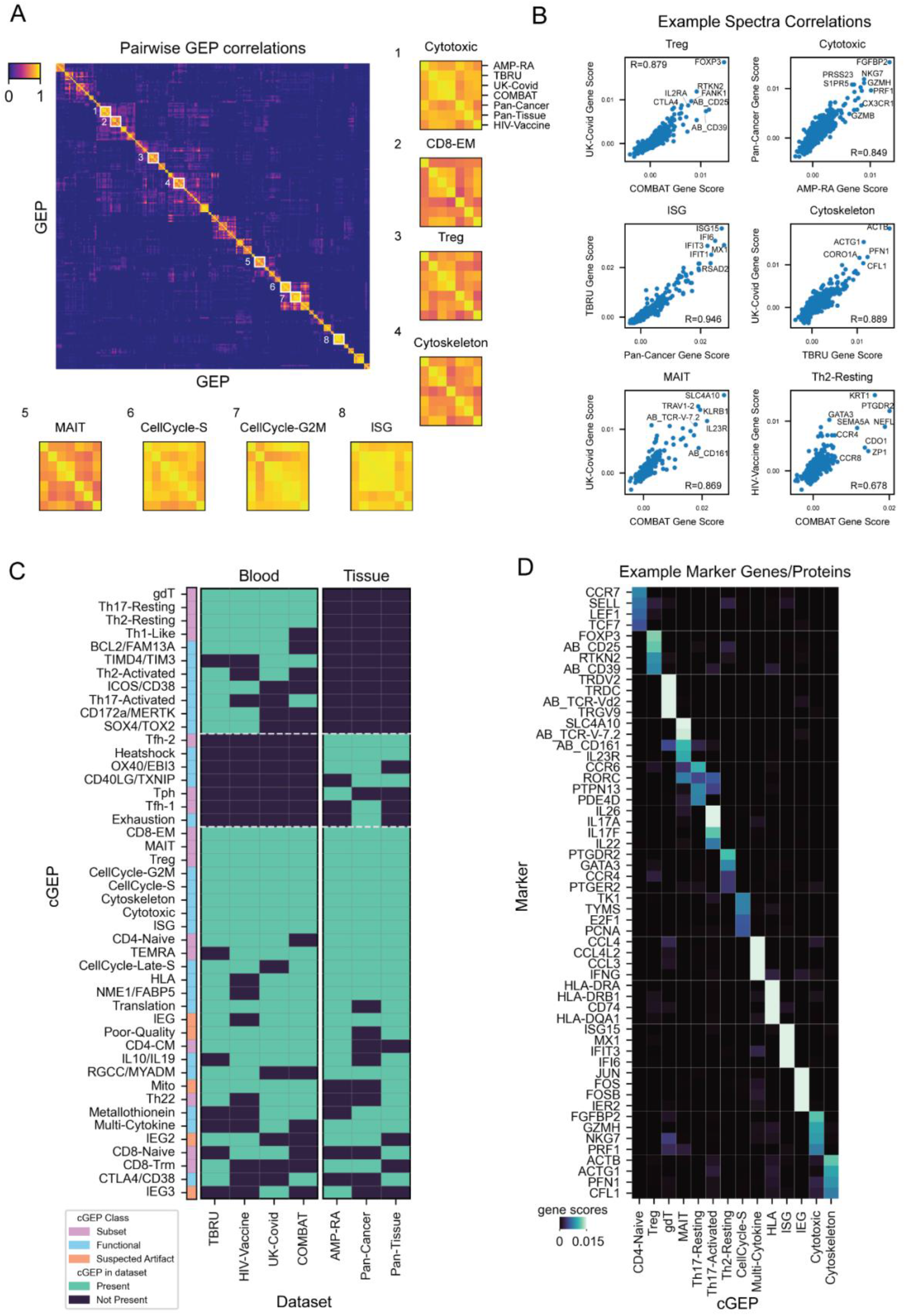
Cataloging consensus gene expression programs (GEPs) across datasets. (A) Pairwise correlations of GEPs discovered across reference datasets with insets for consensus GEPs derived from all seven references. Inset row and column orders are the same for all cGEPs. (B) Scatter plots of selected correlated GEP pairs. X and Y axis labels indicate the datasets the GEP was found in (P<1×10^−100^ for all correlations). (C) Heatmap of cGEPs (rows) and which datasets the comprising GEPs were found in (columns). Green boxes indicate a GEP was found in a dataset. Colorbar indicates the cGEP’s assigned class. cGEPs corresponding to non T-cell lineages were excluded. (D) Marker genes for selected example cGEPs in cNMF gene score units.

We curated a catalog of 46 cGEPs capturing diverse T-cell states, including 11 discovered only in blood datasets, seven discovered only in tissue datasets, and 28 discovered in both (**Table S1**, **Figure 2C**). This represents between 27 and 36 more programs than previous factorization analyses of T-cells^13,15,16^. Of these cGEPs, 43 derived from multiple datasets, while three were singletons found in a single dataset. We excluded 49 of the 52 initially identified singletons since they likely reflect dataset-specific artifacts. The three retained singletons capture disease-or tissue-specific GEPs with a biological justification. For example, the rheumatoid arthritis dataset (referred to as AMP-RA), included a GEP highly enriched for T peripheral helper cells markers (including PD-1 and CD4 protein, *LAG3*, and *CXCL13* RNA), which is characteristic of inflamed rheumatoid arthritis synovium^22^ (**Table S2**). Similarly, the pan-cancer dataset included a cancer-specific exhaustion GEP (*HAVCR2*, *ENTPD1*, *LAG3*) which may be especially enriched in cancer, and a GEP bearing markers for T follicular helper cells (PD-1 protein and *CXCR5*, *IL6R*, and *CXCL13* RNA) which was distinct from a second Tfh-like GEP discovered in multiple non-cancer tissue datasets. In addition to the main T-cell cGEPs, we identified six cGEPs corresponding to non T-cell populations including erythrocytes (*HBA2*, *HBA1*, *HBB*) and plasmablasts (*JCHAIN*, *IGKC*, *IGKV3-20*), potentially derived from doublets. We retained these cGEPs to flag doublet-associated transcriptional signals.

To label cGEPs, we first examined their top weighted genes (**Figure 2D, Supplementary item 2, Table S1-2**). For example, the top 10 weighted genes in the Treg and Th2-Resting cGEPs included the master regulators, *FOXP3* and *GATA3*, respectively. Similarly, top weighted genes helped identify the Th2-Activated (*GATA3, IL4, IL5*) and Th17-Activated (*IL26*, *IL17A*, and *RORC*) cGEPs. Many functional cGEPs could also be readily identified, such as Heatshock (*HSPA1A*, *HSP90AA1*, *HSPA1B*), HLA (*HLA-DRA*, *HLA-DRB1*, *CD74*), Metallothionein (*MT1X*, *MT2A*, *MT1E*), and Actin Cytoskeleton (*ACTB*, *ACTG1*, *PFN1*) (**Figure 2D**).

We also labeled cGEPs based on their ability to discriminate canonical T-cell subsets defined by manual gating on surface markers. We gated PBMC-derived T-cells from the COMBAT CITE-Seq reference dataset^18^ and then used multivariate logistic regression to associate cGEPs with subsets (**Figure S2A**, **Methods**). cGEPs labeled as regulatory T (Treg), gamma-delta T (gdT), mucosal associated invariant T (MAIT), CD4 Naive, CD8 Naive, CD8 effector memory (CD8 EM), CD4 central memory (CD4 CM), and T Effector Memory-Expressing CD45RA (TEMRA) were strongly associated with the expected manually gated populations (P-value<1×10^−200^, Coefficient>0.35, **Figure S2B**). The CD4 effector memory gated population was most strongly associated with cGEPs reflecting expected T-helper subsets labeled as Th17-Resting (*CCR6, RORC, AQP3*) and Th1-like (*IFNG-AS1*, *CXCR3*, and CD195 protein) (P<1×10^−200^ and P=4.1×10^−190^, coefficients 0.36 and 0.22, respectively, **Figure S2B**). Overall, this approach enabled identification of 17 subset-associated cGEPs (**Figure 2C, Table S1**).

As a third strategy to label cGEPs, we used gene-set enrichment analysis with gene-sets from the gene ontology database^23^ and from T-cell polarization experiments^24^ (**Methods, Table S3**). We found that the Th2-Resting and Th2-Activated cGEPs were the most significantly enriched for genes upregulated following 16 hour stimulations of naive T-cells with Th2 polarizing cytokines (Fisher Exact Test OR=22.7, 16.2, P=4.9×10^−5^, 1.7×10^−4^, respectively). Gene set analysis also helped annotate 5 cGEPs corresponding to non-T-cell specific cellular functions including early and late cell cycle S-phase (P=3×10^−56^ for DNA_REPLICATION and P=2×10^−55^ for MITOTIC_CELL_CYCLE), G2M-phase (P=9×10^−74^ CELL DIVISION), interferon stimulated genes (P=1×10^−59^ for RESPONSE TO VIRUS), and translation (P=4×10^−163^ for GOCC_CYTOSOLIC_RIBOSOME).

Next, we identified technical artifact-associated cGEPs that correlate with low-quality cell features (**Table S4**). A cGEP we label Mitochondria contains top markers that are exclusively mitochondrially transcribed genes, which are frequently used to identify low-quality cells^25,26^; as expected, this cGEP had a high correlation with the percentage of mitochondrial reads per cell (average R=0.81 across datasets). We labeled another cGEP Poor-Quality based on its top marker gene *MALAT1*, a long non-coding RNA linked to poor cell viability^27^; this cGEP also correlated with the percentage of mitochondrial transcripts per cell (R=0.25 averaged across datasets, **Figure S2C**) and was inversely correlated with the percentage of protein-coding transcripts per cell (**Figure S2D,** average R=-0.50 across datasets). For the AMP-RA dataset, we had access to raw sequence alignment files so we could quantify the percentage of reads aligned to intergenic regions of the genome; the Poor-Quality cGEP was by far the most correlated with the percentage of intergenic reads per cell (R=0.74, **Figure S2E**). Its usage may be driven by higher levels of contaminating DNA or nascent RNA.

Finally, we label three correlated cGEPs as immediate early gene programs (IEG1, IEG2, IEG3, pairwise R of 0.45-0.70). The top genes include canonical IEGs including *FOS*, *JUN*, and *ZFP36,* and these cGEPs were all enriched for a published IEG gene set^28^ (Fisher Exact Test P<1×10^−53^). We suspect that IEG1 represents the core pathway as it was found in 6 out of 7 datasets (**Figure 2C**) whereas IEG2 and IEG3 represent mixtures with delayed immediate and secondary response genes. We hypothesize that these cGEPs reflect sample processing artifacts in scRNA-Seq, since IEGs are induced in as few as 30 minutes^29^ in response to mitogens or cell stress^30^, and following processing steps like tissue dissociation^31,32^. As evidence of the potential technical nature of these cGEPs, we calculated their mean usage per sample in T-cells, B-cells, NK-cells and monocytes/DCs in the 3 PBMC references. We found that their average usage in T-cells correlates with their usage in other cell-types (R=0.46-0.99, average 0.77, **Figure S2F, Supplementary item 3**), suggesting that they are a sample-intrinsic property, which would be expected of a sample-processing effect. However, in certain contexts, these cGEPs may be biologically important.

### 3. Benchmarking TCAT on an independent query dataset

Next, we benchmarked TCAT on predicting T-cell subsets in an independent CITE-seq dataset. We analyzed 336,739 T-cells from PBMCs of 24 Covid-19-recovered and 17 healthy individuals after flu vaccination^33^ (**Figure 3A**). As ground truth, we assigned cells to one of ten subsets through manual gating of surface proteins (**Figure S3A)**. We then predicted each subset by thresholding the corresponding subset-associated cGEP (**Methods**). For all 10 subsets, thresholding the single most-associated cGEP was comparable to RNA-based hard clustering, across nine different clustering resolutions. Averaged across subsets, the accuracy difference between TCAT and clustering ranged from 0.064 to –0.007 depending on the clustering resolution (**Figure 3B-D**).

**Figure 3.**
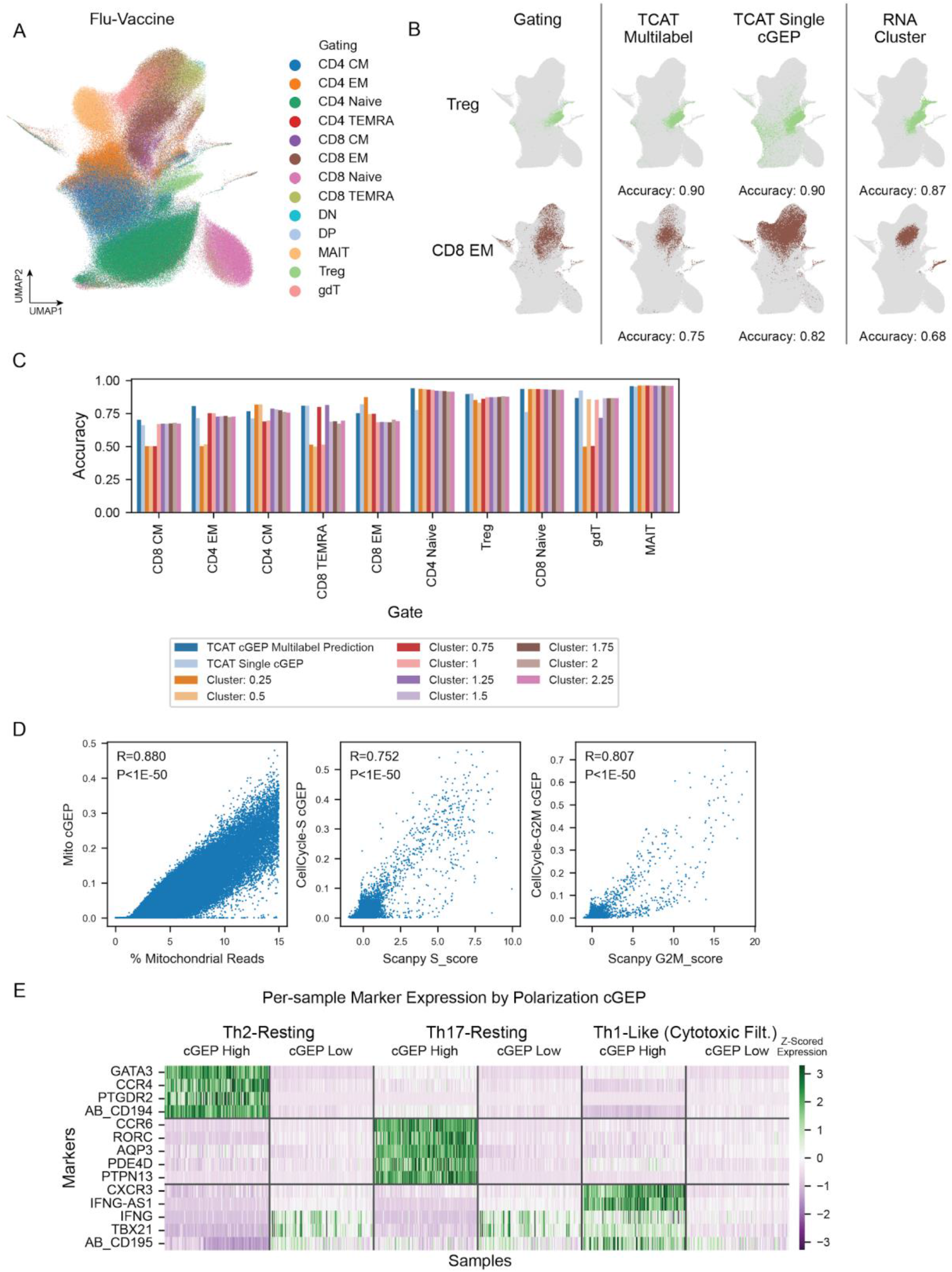
Benchmarking T-CellAnnoTator on a query dataset. (A) UMAP of the Flu-Vaccine dataset colored by the manually gating shown in **Figure S4A**. (B) Same UMAP as (A) but demonstrating prediction of manual gating of Treg and CD8 EM populations with the most associated individual cGEP (usage > 0.025), the multilabel classifier based on multiple cGEPs, or Ledien clustering with resolution 1.0. (C) Comparison of balanced accuracy for prediction of manually gated subsets, including clustering with multiple Leiden resolution parameters. (D) Usage of the mitochondria cGEP against the percentage of mitochondrial reads per cell (left). Usage of the CellCycle-S (middle) and CellCycle-G2M (right) cGEPs against the S and G2M scores output by Scanpy’s score_genes_cell_cycle function with published proliferation gene sets^34^. (E) Heatmap of pseudobulk expression in cGEP-high and low cells, per sample. Samples are normalized by library size and expression is z-scored across rows.

Since subsets can contain heterogeneity not captured in univariate analysis (e.g. multiple polarized populations within CD4 effector memory), we performed multivariate analysis using all cGEPs for simultaneous multi-label prediction (**Methods**). We trained the classifier on the COMBAT dataset and evaluated its performance on the Flu-Vaccine dataset. The classifier was more accurate than RNA clustering across all nine clustering resolutions tested, with average accuracy differences ranging from 0.10 to 0.033 (**Figure 3B-C, Figure S3B-C**). Thus, for PBMC-derived T-cells, TCAT can be combined with a multilabel classifier to predict subsets without requiring manual annotation.

We also compared TCAT’s subset classification accuracy against NMFproject^13^ and gene-sets derived from a recent NMF analysis of tumor-infiltrating T-cells^16^ (**Methods**). TCAT single cGEP and multi-label classification yielded higher area under the curve (AUC) for all lineage predictions than these other approaches (**Figure S3B-C)**.

Next, we validated TCAT’s prediction of functional cGEPs relative to common continuous metrics. Usage of the mitochondrial cGEP was highly correlated with percentage of mitochondrial reads (R = 0.88, **Figure 3D**). In addition, predicted cell cycle cGEP usages corresponding to the S and G2M phase were highly correlated with cell cycle scores calculated from corresponding published gene sets^34,35^ (R=0.75-0.81, **Figure 3D**).

Finally, we validated prediction of T-cell polarization against expression of canonical markers. We discretized cells based on their expression of the Th1-Like, Th2-Resting, and Th17-Resting cGEPs (usage>0.1) and computed per-sample pseudobulk profiles of high and low usage cells. Th2-Resting-high samples expressed significantly more *GATA3*, *CCR4*, and *PTGDR2* than Th2-Resting-low samples (P<1×10^−35^ all, paired T-test) (**Figure 3E**). Th17-Resting-high samples also had increased expression of Th17 markers including *CCR6*, *RORC*, and *AQP3* (P<1×10^−55^ all). The Th1-Like-high samples had increased expression of the Th1 markers *CXCR3*, *IFNG-AS1*, and CD195 protein (P<1×10^−35^ all). However, the Th1 markers *IFNG* and *TBX21* were also expressed in Th1-Like-low samples (**Figure S3D**). We suspected this was due to the known expression of these genes in cytotoxic T-cells^36,37^. When we excluded cells high in the cytotoxic cGEP (usage>0.1) prior to pseudobulking, *IFNG* and *TBX21* were significantly higher in Th1-Like-high samples (P=8.2×10^−13^, P=9.6×10^−47^, **Figure 3E**, **S3D**). Thus, TCAT can predict T-cell polarization in query datasets.

### 4. cGEPs capture multi-program identities of T-cells in scRNA-Seq

Next, we illustrate how TCAT can reveal cellular heterogeneity not visible with clustering. Using the COMBAT dataset as an example, we analyzed cell cycle, a common signature that frequently obscures other aspects of proliferating cells^38^. In the initial publication, two clusters were annotated as proliferating CD4s and CD8s with subclusters that didn’t clearly correspond to subsets (e.g. CD4.TEFF.prolif.1, CD4.TEFF.prolif.GZMB.1). One sub-cluster labeled CD4.TEFF.prolif.MKI67lo was enriched for the myeloid doublet cGEP (**Figure 4A-B**) and expressed myeloid marker genes (e.g. *CD14*, *MNDA*, **Supplementary item 4**), illustrating how cell cycle can drive cells with distinct cell lineages to cluster together. By contrast, TCAT readily identified distinct proliferating subsets based on co-expression of cell cycle and subset cGEPs, including CD8 EMs, TEMRAs, and Treg (**Figure 4C-D**).

**Figure 4.**
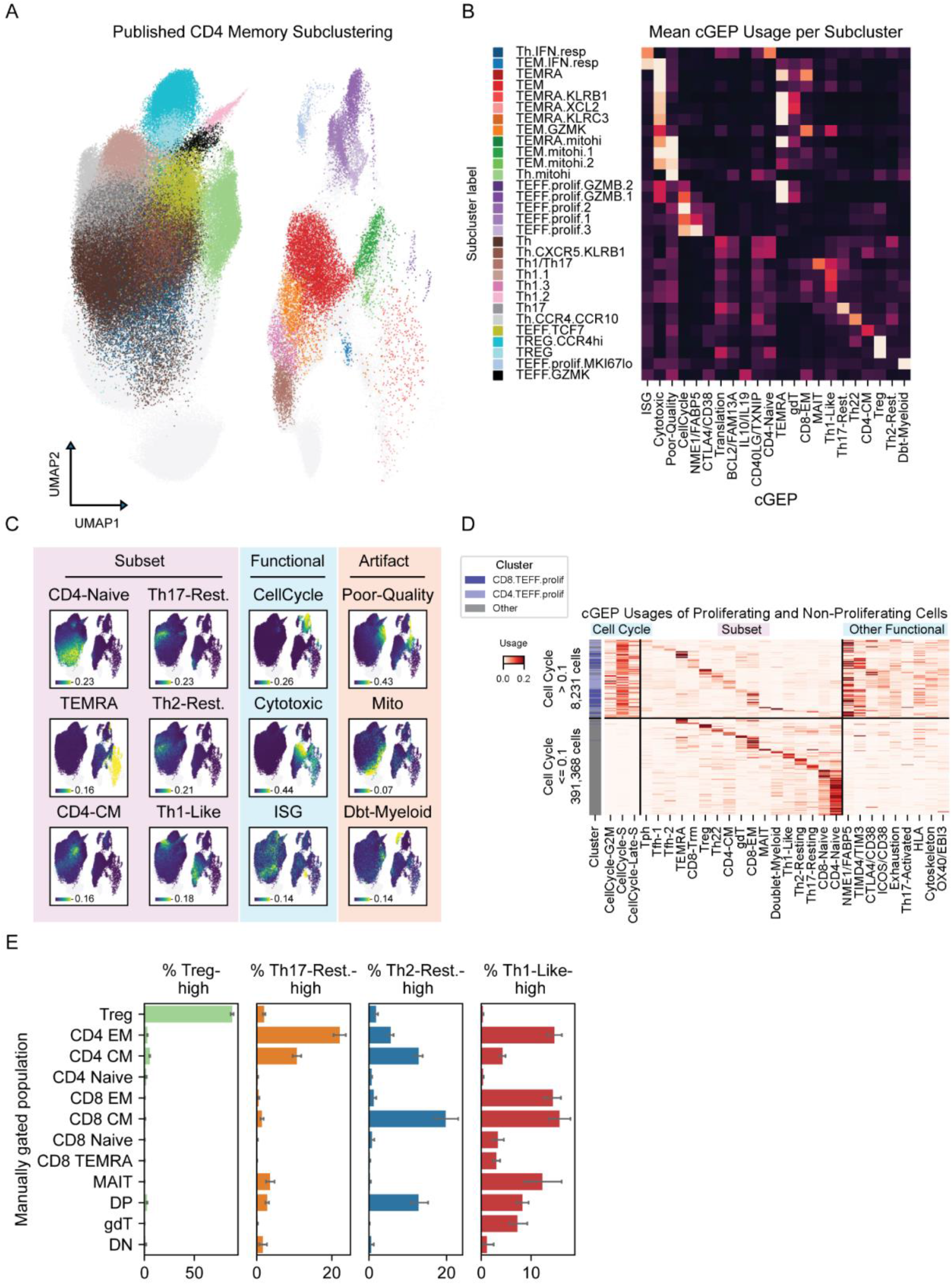
Comparing TCAT to clustering in the COMBAT dataset. (A) UMAP of T-cells showing published sub-clusters of clusters annotated as CD4 memory with other clusters shown in gray. (B) Average usage of selected cGEPs across CD4 memory subclusters. (C) Same UMAP as (A) but colored by usage of selected subset, functional, and artifact cGEPs usage. Intensities are averaged over 20 nearest neighbors to reduce overplotting. (D) Usage of selected cGEPs in cells with high or low usage of cell cycle GEPs. Cells are grouped by their most highly used subset GEPs.(E) Percentage of cells within each manual gate assigned to each polarization (usage > 0.1). Bar represents the average and whiskers represents the 95% confidence interval, across samples.

Disentangling cell cycle and subset enabled us to quantify the percentage of proliferating cells per subset and disease status. We assigned cells to subsets based on their most highly used subset cGEP. This revealed increased expression of cell cycle cGEPs across many T-cell subsets in Covid-19 compared to healthy cells, in both Covid-19 datasets (**Figure S4A**). The most proliferative subsets in both Covid-19 and control samples expressed the T peripheral helper cGEP, reflecting an inflammatory population that was recently identified in Covid-19^39^.

We identified other functional cGEPs that obscured T-cell subsets, akin to proliferation. Many CD4 memory subclusters in the original study were most strongly enriched for functional cGEPs such as ISG, Cytotoxicity, and Poor-Quality, rather than subset cGEPs (**Figure 4B-D, Supplementary item 4**). The CD4.Th.mitohi and CD4.Tem.mitohi.1 clusters were driven by high usage of the Poor-Quality cGEP and contained cells expressing multiple subset cGEPs. The CD4.TEM.IFN.resp and CD4.Th.IFN.resp clusters were both predominantly driven by the interferon stimulated gene (ISG) cGEP. The CD4.TEM.IFN.resp cluster had high usage of the Cytotoxicity and TEMRA cGEPs while the CD4.Th.IFN.resp cluster contained cells expressing many subset cGEPs including CD4-Naive (**Figure 4B, S4B**). Cells with high usage of the CD4-Naive cGEP expressed CD4 naive markers including CD45RA protein and *SELL* RNA, confirming that clustering had misclassified them as memory T-cells (**Supplementary item 4**).

Clustering also obscured the subset of CD4 T-cells expressing the Cytotoxicity cGEP. We visualized the per-cell usage of all cGEPs in cells from the CD4 memory sub-clusters that had high Cytotoxicity cGEP usage (average cluster usage>0.1, **Figure 4B**). Intriguingly, these clusters contained heterogeneous cells with high usage of many subset cGEPs including CD8-EM, Th1-Like, TEMRA, and gdT (**Figure S4C**). Pseudobulk analyses showed that cells co-expressing these cGEPs (usage>0.1 for both) co-expressed the expected cytotoxicity and subset marker genes (**Figure S4D**). Thus, TCAT can reveal subset heterogeneity within cytotoxic T-cells.

TCAT could readily annotate polarization status based on usage of the Th1-Like, Th2-Resting, and Th17-Resting cGEPs (**Figure 4C**). By contrast, the published clustering did not identify a Th2 cluster, and clusters annotated as Th1 and Th17 were only identified with a high clustering resolution resulting in 243 clusters, likely due to other conflating signals. As expected, there was significant enrichment between cells annotated as Th1 by clustering and high Th1-Like cGEP usage, as well as Th17 clustering and high Th17-Resting cGEP usage (P<1×10^−100^ for both, fisher exact test).

However, TCAT additionally identified expression of polarization cGEPs outside of the CD4 memory compartment (**Figure 4E**). We annotated polarization across manually gated T-cell subsets with a usage threshold>0.1. As a control, we confirmed that the Treg cGEP was highly enriched in the Treg gate, with an average of 88.1% of gated Tregs expressing the cGEP, compared to 5.3% for the next highest population. Similarly, the Th17-Resting cGEP was most enriched in the expected CD4 EM (22.1%) and CD4 CM (10.7%) populations compared to only 3.5% for MAITs, the next highest. Surprisingly, the Th2-Resting cGEP was most commonly assigned within the CD8 CM (19.8%), CD4 CM (12.8%), and CD4/CD8 Double Positive (12.8%) populations. The Th1-Like cGEP was also used by CD8 T-cells; it was most prevalent within the CD8 CM (15.7%), CD4 EM (14.7%), CD8 EM (14.4%), and MAIT populations (12.3%). The calculated subset polarization proportions were highly correlated between the COMBAT and Flu-Vaccine datasets, the two datasets with the best quality manual gating (R>0.9, P<5.5×10^−5^ for all three, **Figure S4E**). Furthermore, cells assigned to each polarization had high usage of the expected marker genes for that polarization, irrespective of whether they were CD4+ or CD8+ (**Figure S4F**). These findings support the emerging recognition of polarized CD8 T-cell populations^40^ and illustrate how these populations are easily revealed by TCAT.

### 5. cGEPs associated with TCR-dependent activation

Next we identified cGEPs induced following antigen recognition by the TCR. To do so, we developed AIM-Seq (Activation-Induced Marker (AIM) assay followed by scRNA-Seq), an assay to profile T-cells after antigen stimulus (**Figure 5A-D**). We collected PBMCs from 5 genome-wide genotyped healthy donors and stimulated them for 24 hours using a pool of 176 peptide antigens from common pathogens (CEFX, JPT)^41^ and anti-CD28/CD49d co-stimulation. Using flow cytometry, we separated T-cells expressing activation-induced markers (OX40 and PD-L1 for CD4s^42^, CD137 for CD8s^43^, AIM-positive) from unactivated cells (negative for these markers, AIM-negative). As a negative control, we activated cells non-specifically with anti-CD28/CD49d costimulation without peptides (Mock). We labeled cells from these conditions with hashtag antibodies and pooled them for single-cell RNA, CITE, and TCR repertoire sequencing (**Methods**).

**Figure 5.**
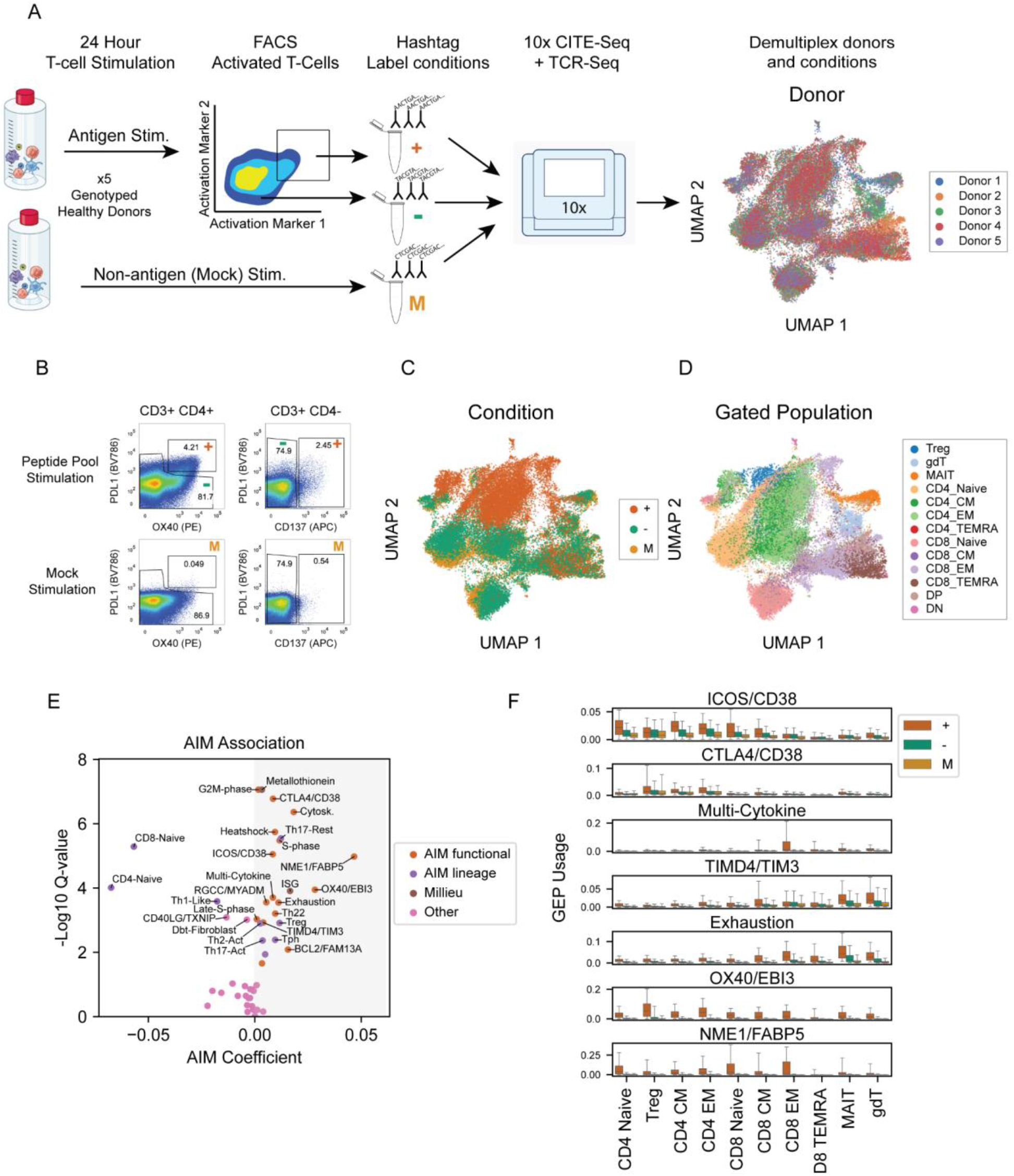
Identifying cGEPs associated with TCR-dependent activation. (A) Schematic of AIM-Seq. (B) FACS experiment from an AIM-Seq run showing surface activation markers in CD3+CD4+ and CD3+CD4-gated populations with the gates used for AIM-positive (+), AIM-negative (-) and Mock (M) populations. (C-D) UMAP of AIM-Seq dataset colored by sorting condition (C) or manually gated population (D). (E) cGEP association with AIM-positive samples. X-axis shows the mean Log_2_ ratio of average usages. Y-axis shows the –Log_10_ P-value. cGEPs are labeled by assigned category. (F) Average usage of selected Aim-associated cGEPs in +, –, and U cells from different gated subsets. Boxes represent interquartile range. Error bars represent 95th percentiles.

As expected, CEFX stimulated CD4 and CD3+CD4-(hereafter labeled CD8) T-cells contained higher proportions of AIM-positive cells than mock (**Figure 5B, S5A**). 4.21% of CD4 T-cells and 2.45% of CD8s were AIM-positive, compared to 0.049% and 0.54% of mock-stimulated CD4 and CD8 T-cells, respectively.

The CITE-Seq data showed that AIM-positive cells expressed additional surface activation markers including CD54, CD25, CD71, and CD69 beyond the sorting markers (T-test P<1×10^−200^, **Figure S5C-E**). Moreover, AIM-positive cells were significantly depleted of naive T-cells (P= 0.027 and P=8.6×10^−4^, for CD4 and CD8, respectively) and enriched for Tregs, CD4 central and effector memory populations (P =0.00064, 0.0044 and 0.054, respectively, **Figure S5F**). This is unsurprising as the peptide pool is derived from common pathogens and prior memory is expected. However, 11.8% of the AIM-positive cells were CD4 naive and 1.4% were CD8 naive, indicating we could detect both memory and naive cell responses.

Next, we identified cGEPs associated with antigen-specific activation in this assay. We used pseudobulk sample-level regression to identify cGEPs upregulated in AIM-positive cells relative to AIM-negatives (**Methods**). This identified 24 significant positively associated cGEPs (false discovery rate (FDR) corrected P < 0.05), including two that are milieu regulated (I.e. non TCR-dependent), five representing enriched subsets, and 17 functional cGEPs (**Figure 5E, S5G**).

The two milieu mediated cGEPs, Interferon Stimulated Gene (ISG) and Metallothionein, were significantly upregulated in both AIM-negative and AIM-positive cells relative to mock (ISG: AIM-negative – P=8.9×10^−7^, AIM-positive – P=3.1×10^−5^; Metallothionein: AIM-negative – P=1.5×10^−3^, AIM-positive – P=3.3×10^−9^). Interferon is a secreted cytokine that can activate nearby cells independent of TCR-activation to induce the ISG cGEP. Shifting extracellular cytokine or ion concentrations may similarly induce TCR-independent upregulation of the metallothionein cGEP^44^.

Five subset-associated cGEPs were increased in AIM-positive cells relative to AIM-negatives (Th17-Resting, Treg, Tph, Th22, and Tfh-2) and 3 were increased in AIM-negatives (CD8-Naive, CD4-Naive, and Th1-like) (**Table S5**). These associations likely reflect differential abundance of cell populations rather than upregulation of the cGEPs, consistent with the manual gating results (**Figure S5E**).

The remaining 17 AIM-associated programs are functional cGEPs including many with well-known links to TCR-stimulation. Six of these are not T-cell specific, namely the three cell cycle cGEPs^45^ (P<3.6×10^−4^), actin cytoskeleton^46^ (P=3.3×10^−8^), heatshock^47,48^ (P=1.7×10^−7^), and MHC class II^49^ (P=0.012).

Excluding these leaves 11 functional AIM-associated cGEPs that may be specific to T-cell activation. These include CTLA4/CD38 (P=9.7×10^−9^), ICOS/CD38 (P=1.5×10^−6^), NME1/FABP5 (P=2.0×10^−6^), OX40/EBI3 (P=2.6×10^−5^), Multi-cytokine (P=5.4×10^−5^), Exhaustion (P=9.3×10^−5^), TIMD4/TIM3 (P=5.0×10^−4^), Th2-Activated (P=5.9×10^−4^), Th17-Activated (P=2.1×10^−3^) and BCL2/FAM13A (P=4.3×10^−3^). We highlight 4 of these cGEPs here. CTLA4/CD38 showed the most upregulation in Tregs and CD4 memory cells (**Figure 5F**) and is characterized by CD278 and CD38 protein levels as well as the anti-inflammatory genes *CTLA4* and *IL10*. ICOS/CD38 has similar top markers including CD278, CD71, and CD38 but shows broad upregulation across naive T-cells and CD4 memory cells. The OX40/EBI3 cGEP includes many of the activation-induced markers used to define AIM positivity in the first place including *TNFRSF4* which encodes OX40 and *IL2RA* which encodes CD25. TIMD4/TIM3 is most expressed in MAIT, gdT, and CD8 memory T-cells and is characterized by expression of activation markers (CD38 protein and RNA) and cytotoxicity genes (*GZMB*, *GZMA*, *GNLY*), and likely represents a cytotoxic activation response.

We hypothesized that AIM-associated cGEPs would be enriched in proliferating cells *in vivo* since proliferation is a core response to TCR activation. To test this, we performed pseudobulk sample-level association tests to identify cGEPs with higher usage in proliferating cells (sum of cell cycle cGEPs>0.1) than non-proliferating cells (sum<0.1, **Methods**). The results were highly concordant across datasets (**Table S6, Supplementary item 5**). 15 cGEPs were significantly upregulated with proliferation in at least four out of six datasets. Meta-analysis across datasets identified 12 functional cGEPs (including the three cell cycle cGEPs) and two subset cGEPs (Th17-Activated and Tph) that were significantly associated with proliferation (**Figure S5H**). Consistent with our hypothesis, 14 of 15 proliferation-associated cGEPs (including the 3 cell cycle cGEPs) were upregulated with AIM positivity (Fisher exact test P=2.1×10^−5^). Thus, the AIM-associated cGEPs are associated with proliferation *in vivo*, consistent with a role downstream of TCR activation.

### 6. Annotating antigen-dependent activation *in vivo*

Next, we developed a per-cell antigen-specific activation (ASA) score to identify and characterize TCR-activated T-cells in disease. We used forward stepwise selection to select AIM-associated cGEPs that predicted co-expression of the activation markers CD71 and CD95 in the COMBAT and Flu-Vaccine datasets (**Methods**). These markers show sustained upregulation within less than 24 hours of TCR activation^50–53^, were upregulated in the AIM-positive cells (**Figure S5D-F**), and had high quality across subsets in both datasets (**Figure S6A**). Stepwise optimization defined ASA as the sum of four cGEPs – TIMD4/TIM3, ICOS/CD38, CTLA4/CD38, and OX40/EBI3 (**Figure S6B**, **Methods**).

ASA accurately classified T-cells with CD71/CD95 co-expression suggestive of TCR-activation, yielding AUCs of 0.920 and 0.818 in the COMBAT and Flu-Vaccine datasets (**Figure S6C-D**). It also predicted AIM positivity with an AUC of 0.828 in the AIM-Seq assay (**Figure S6E**) and was correlated with other surface markers of activation (e.g. R=0.43 (CD69) and 0.52 (CD25), P<1×10^−100^, **Supplementary item 6**). For cases where a discrete label is preferable to a continuous score, we picked an ASA threshold of 0.0625 based on the trade-off between sensitivity and specificity (**Figure S6C-E**). With this threshold, ASA annotated 76.7% of CD71+CD95+ and 5.2% of non-CD71/CD95 double positive T-cells in the COMBAT dataset (**Figure 6A**). In the AIM-Seq dataset, ASA annotated 60.6%, 7.0%, and 3.2% of stimulated AIM-positive, stimulated AIM-negative, and mock stimulated cells, respectively (**Figure 6B**).

**Figure 6.**
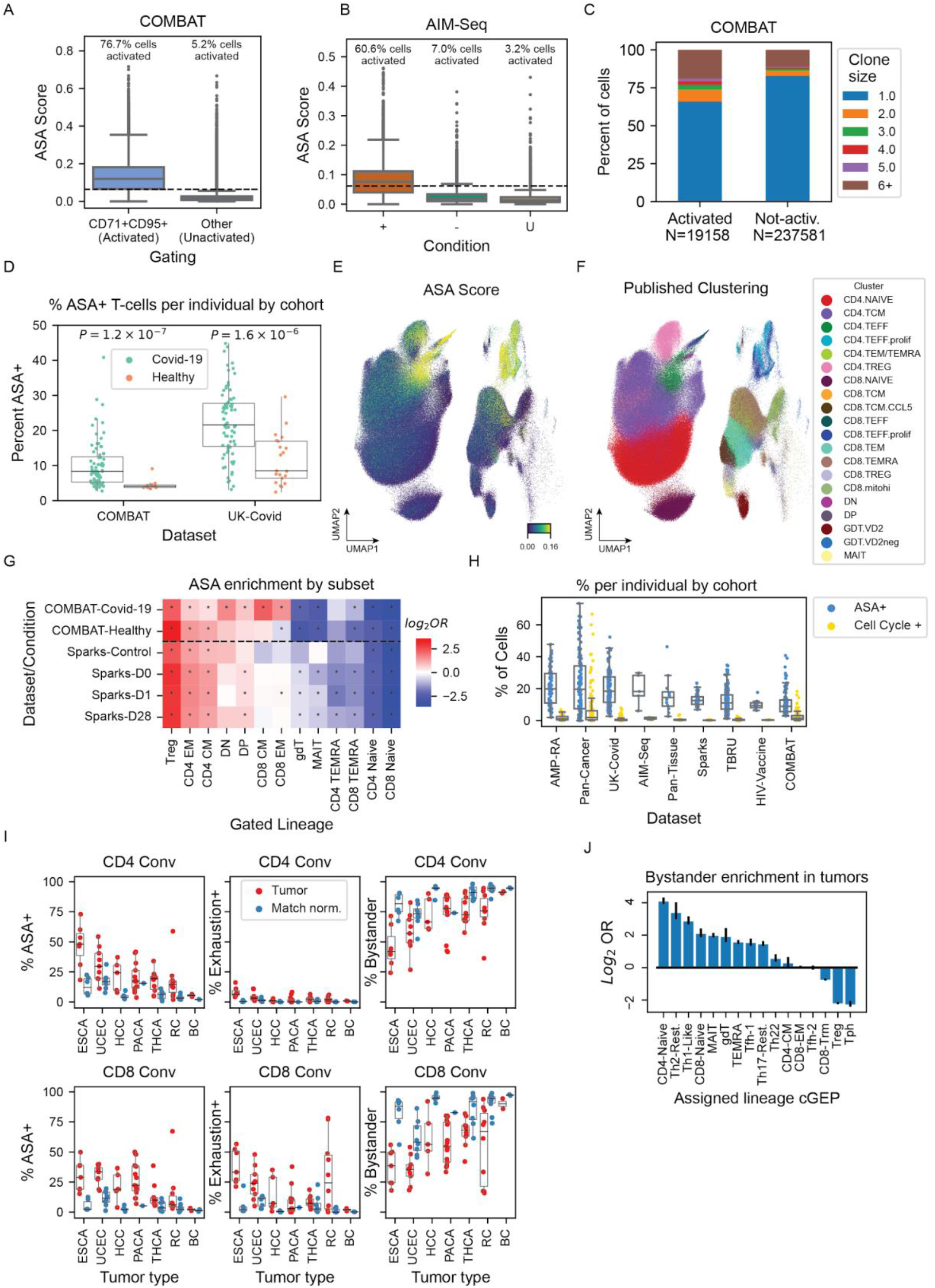
Annotating antigen-specific activation (ASA) *in vivo*. (A) Box plot of ASA score for cells stratified as activated (CD71+CD95+) or not activated. (B) Same as (A) but for AIM-Seq with cells stratified by sort condition. (C) Clonality in manually gated conventional CD4 and CD8 T-cells annotated as activated (ASA>0.065) or not activated (ASA<0.065). Clonality is defined as the number of cells in the same sample with an identical alpha and beta CDR3 amino acid sequence. (D) Percentage of activated CD4 and CD8 convs (ASA>0.065) in Covid-19 and healthy control samples, by cohort. (E-F) UMAP of the COMBAT dataset colored by ASA score or low-resolution published clustering. (G) Log_10_ odds ratio for 2×2 association of ASA positivity and manual gating subset assignment. * indicates P-value<0.05. (H) Percentage of activated (ASA>0.065) or proliferating (sum of cell cycle cGEPs>0.1) cells per sample across datasets. Boxes represent the interquartile range and whiskers represent 95% quantile range. (I) Percentage of activated, exhausted (exhaustion cGEP usage>0.065), or bystander (ASA + exhaustion usage<0.065) T-cells in CD4 and CD8 Convs, per sample stratified by tumor type and corresponding healthy tissues. (J) Log_2_ odds ratio for enrichment of bystander T-cells by subset cGEP assignment. Error bars represent 95% confidence intervals.

As proliferation is a core response to activation, we found high ASA in proliferating T-cell clusters (**Figure 6E-F**) and significant overlap of ASA-high and proliferating cells (specifically, cells with summed cell cycle usage > 0.1, Fisher Exact OR 2.8-58.8, P<1×10^−100^, **Figure S6F – left**). However, across reference datasets, substantially more cells were annotated as ASA-high than proliferating (P=8.8×10^−189^, paired T-test, **Figure 6H**). Consistent with this, correlation between summed cell cycle cGEP usage and ASA was relatively low (mean=0.15) (**Figure S6F – right**). Thus, while proliferation and antigen-specific activation overlap to some extent, ASA offers greater sensitivity for classifying TCR-activation.

As clonal expansion often follows TCR activation, we tested whether high clonality was associated with ASA in Covid-19 patients. ASA-high cells were more likely to be clonal, i.e. have a TCR found in multiple cells from the same sample (Fisher Exact Test: COMBAT OR=2.50, UK-Covid OR=2.28, P< 1×10^−100^ for both). Binarized ASA and cell cycle status were independently associated with clonality in a multivariate logistic regression (ASA Beta = 0.45, 0.50; Cell cycle Beta = 0.66, 0.52 in COMBAT and UK-Covid respectively, P<1×10^−22^, **Methods**). Furthermore, the absolute number of cells sharing a TCR sequence in a sample was significantly higher in ASA-high than ASA-low cells (Mann Whitney U test P<1×10^−100^, both datasets, **Figure 6C, S6G**).

Next, we evaluated how ASA varied between Covid-19 and healthy samples across T-cell subsets. The percentage of activated (I.e. ASA positive) conventional T-cells varied widely across samples, between 2.7%-41.2% (mean 10.3%) and 4.9%-44.7% (mean 22.1%), in the COMBAT and UK-Covid datasets, respectively (**Figure 6D**). Activation rates were significantly higher in conventional T-cells in Covid-19 samples than in healthy controls (COMBAT P=1.9×10^−7^, UK-Covid P=1.5×10^−6^), even in CD4+ and CD8+ T-cells separately (**Figure S6H-J**). Activation rates were similar between CD4s and CD8s (median activation of 8.3%, 21.8% for CD4s and 7.8%, 21.7% for CD8s in COMBAT and UK-Covid). By contrast, there was greater Treg activation in both healthy and Covid-19 samples, with a median of 33.6 and 35.3% of cells activated in COMBAT and UK-Covid (**Figure S6J**). This coincided with substantial overlap of ASA with the Treg cluster (**Figure 6E-F**). Tregs were the most ASA-enriched subset in healthy control samples in the COMBAT (OR=11.4, P<1×10^−100^) and Flu-Vaccine datasets (OR=4.1, P<1×10^−10^) (**Figure 6G**). Outside of acute infection, we would expect Tregs to be actively suppressing inappropriate activation. By contrast, in acute Covid-19 samples, we saw less enrichment for Tregs (OR=4.8 down from 11.4) and more for CD8 central memory (OR=4.8), CD8 effector memory (OR=2.8), and double negative populations (OR=3.1), reflecting the antiviral response (all P<1×10^−10^).

Next, we quantified levels of T-cell exhaustion and activation per sample and subset within the pan-cancer dataset. CD4 conventional T-cell (CD4 Conv) activation rates varied widely across and between tumor types (**Figure 6I)**. The highest rates of activation were in esophageal cancer (ESCA – median 48.0%) and the lowest were in bladder cancer (BC – median 5.4%, **Figure 6I – left**). As expected, there was minimal exhaustion usage by CD4 Convs across cancer types^54^ but highly variable levels of CD8 conventional T-cell (CD8 Conv) exhaustion (**Figure 6I – middle**). The percentage of activated CD4 Convs and CD8 Convs was correlated (R=0.70, P=2.6×10^−9^). In addition, CD4 conv activation was somewhat correlated with CD8 Conv exhaustion (R=0.38, P=4.0×10^−3^, **Figure S6K**). CD4 Treg activation levels were higher in healthy tissues and tumors than CD4 and CD8 Conv T-cells (**Figure S6L**). In addition, Treg activation was significantly higher in thyroid cancer (P=3.0×10-6) and esophageal cancer (P=0.0045) relative to matched normal tissues.

Observing that many tumor-infiltrating T-cells had both low ASA and exhaustion usage, we defined bystanders as cells with summed ASA and exhaustion usage below 0.0625. The percentage of CD4 bystanders varied widely by cancer from 42.0% (esophageal) to 91.2% (bladder) and CD8 bystanders varied similarly from 35.5% (endometrial) to 90.1% (bladder).

Within tumor samples, we tested which T-cell subset cGEPs were enriched for bystanders (**Figure 6J**). The most bystander-enriched subsets were CD4-Naive (OR=15.9), Th2-Resting (OR=10.6), Th1-like (OR=7.3), MAIT (OR=4.42), and CD8-Naive (OR=4.03) (Fisher Exact Test P< 1×10^−100^ for all comparisons). The subsets most depleted of bystanders were also those most enriched for activation, namely Tph (OR=0.19), Treg (OR=0.23), and CD8-Trm (OR=0.61) (P<1×10^−21^, all comparisons). These analyses illustrate how TCAT and ASA scoring can facilitate exploration of disease.

### 7. Identifying disease-associated cGEPs

Next, we associated cGEPs with sample-level disease phenotypes in infection, autoimmunity, and cancer (**Table S7**). First, we tested cGEP associations with Covid-19 (**Methods**). We applied ordinary least squares using psuedobulk sample-level features to two PBMC-derived T-cell datasets: UK-Covid (80 Covid-19, 21 healthy donors, **Figure 7A**) and COMBAT (77 Covid-19, 10 healthy donors, **Figure 7B**). We observed overall concordant cGEP associations (Pearson R=0.64, P=2.8×10^−7^, **Figure 7C**). Consistent with the key role of interferon in viral infections^17,18^, ISG was the most positively upregulated cGEP in both datasets (FDR-corrected P, denoted as Q<0.05). AIM-associated functional cGEPs were up-regulated in acute Covid-19, consistent with viral activation of T-cells. These included exhaustion, cell cycle, TIMD4/TIM3, OX40/EBI3, NME1/FABP5, and CTLA4/CD38 (Q<0.05 for both datasets). We also found increased Tph cGEP usage in Covid-19 relative to controls (Q<1×10^−8^ for both datasets), consistent with recent demonstration of increased abundance of this subset in infection^39^. An intriguing novel finding is that the Th1-like cGEP was significantly negatively associated with Covid-19 in both datasets (Q<1×10^−4^). This negative association was seen within manually gated CD4 memory (Q=1.1×10^−4^) and CD4 effector memory subsets (Q=4.5×10^−6^), suggesting it is not due to differential abundance of circulating memory CD4 T-cells. Consistent with this, pseudobulk expression of the Th1 markers *CXCR3* RNA and protein levels were significantly lower in Covid-19 samples relative to controls (P=8.1×10^−7^ and 0.010 respectively, COMBAT). Immediate early gene cGEPs (IEG1, IEG2, IEG3) were also significantly associated with Covid-19 in the COMBAT dataset (FDR-corrected P<1×10^−5^) but not in the UK-Covid dataset (P>0.5), perhaps related to sample processing differences (see section 2).

**Figure 7.**
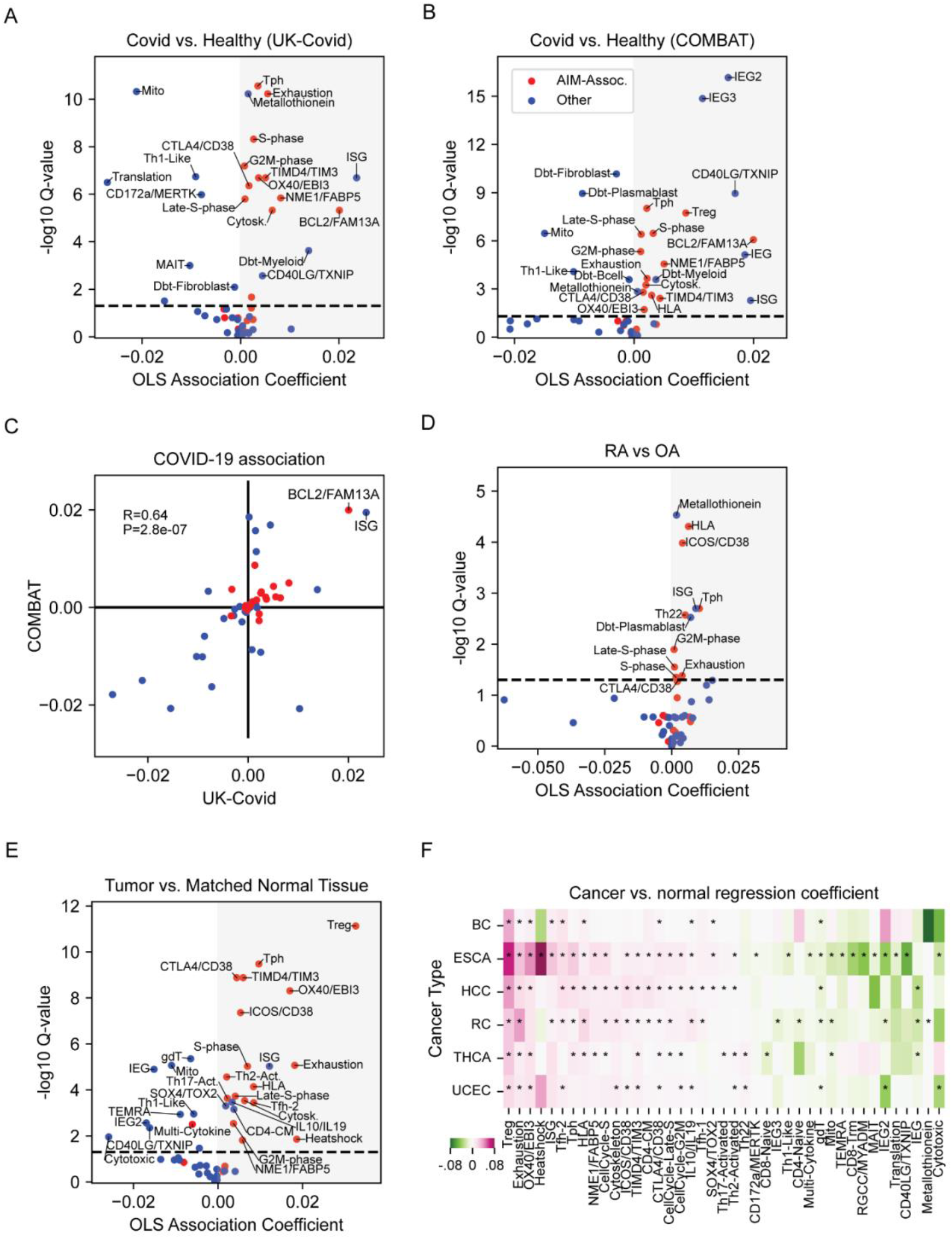
cGEPs association with disease. (A-B) Associations of cGEP usage with Covid-19 status for UK-Covid and COMBAT datasets. X-axis shows the regression coefficient. Y-axis shows the –Log10 FDR-corrected Q-value. (C) Scatter plot of regression coefficients from (A) and (B). (D-E) Same as (A) but comparing synovial T-cells from patients with Rheumatoid Arthritis and Osteoarthritis, or from tumors and healthy adjacent tumors. (F) Regression coefficients for tumor vs. normal samples for each tissue of origin. * denotes P<.05 for the corresponding coefficient. Cancer type abbreviations are: bladder cancer (BC), esophageal cancer (ESCA), hepatocellular carcinoma (HCC), renal cell carcinoma (RC), thyroid carcinoma (THCA), and endometrial cancer (UCEC).

Next, we identified cGEPs associated with inflamed synovial tissue in rheumatoid arthritis (RA) using the AMP-RA dataset, which includes synovial biopsies from 70 RA and 8 osteoarthritis (OA) patients (**Figure 7D**)^20^. Ten out of the eleven significantly associated cGEPs were AIM-associated, including the metallothionein (Q=2.9×10^−5^), ISG (Q=0.0020), Tph (Q=0.0020), HLA (Q=4.9×10^−5^), ICOS/CD38 (Q=0.00010), Exhaustion (Q=0.041), and cell cycle (Q<.05 for all three). Of note, Metallothionein was shown to be increased in the plasma of RA patients and within the synovia of mouse models of RA^55^. The Tph association is consistent with prior observations by us and others of Tph enrichment within RA synovia^22^. The Th22 cGEP was also associated with RA (Q=0.0027), confirming a prior observation of increased Th22 cell abundance in RA synovia, where they may stimulate osteoclasts^56^.

Lastly, we identified cGEPs associated with T-cells in tumors relative to matched healthy tissues (**Figure 7E**). We utilized a pan-cancer dataset containing 89 tumor and 47 matched normal samples from 13 cancer types. First, we analyzed all samples together, controlling for tumor type and sequencing technology as fixed effects. The Treg cGEP was the most strongly associated, consistent with the known importance of Tregs in tumors (Q=7.4×10^−12^)^57^. The exhaustion and ISG cGEPs were also strongly associated with cancer, as expected (Q=8.5×10^−6^ and 9.3×10^−6^, respectively)^58,59^. There was also substantial upregulation of AIM-associated functional cGEPs, including CTLA4/CD38 (Q=1.3×10^−9^), TIMD4/TIM3 (Q=1.3×10^−9^), and OX40/EBI3 (Q=4.9×10^−9^). Overall, 17 of the 21 significantly upregulated cGEPs in tumor-infiltrating T-cells were AIM-associated (Fisher exact test P=7.4×10^−6^).

We also separately tested for cGEP association in each of the six cancer types with at least two normal and two tumor samples (**Methods**). The results were highly concordant across cancers (P<.05, sign test, for 14 out of 15 pairs of tumor types, **Figure 7F**). For example, the Treg, Exhaustion, and CTLA4/CD38 cGEPs were significantly upregulated in all six tumor types tested (P<.05). However, some signals were more specific. The Th17-Activated cGEP was only significant in thyroid and hepatocellular carcinoma (P=5.3×10^−6^ and P=0.013), while the Th2-Activated cGEP was upregulated in esophageal, uterine, thyroid and hepatocellular carcinoma (P=0.023, P=0.023, P=0.00057, P=0.0019).

Surprisingly, the Tfh-2 and Tph cGEPs were both upregulated in cancer (Q=3.6×10^−4^, Q=3.3×10^−10^). T follicular helper (Tfh) and T peripheral helpers (Tph) are *CXCL13*-producing CD4 subsets that recruit B-cells and aid in antibody production. Tfhs are found primarily in lymphoid organs and Tphs are predominantly in inflamed tissues^60^, including likely within tumors^61^.

Consistent with functional Tph activity, the expression of the B-cell chemoattractant *CXCL13* was highly correlated with average Tph cGEP usage across samples (R=0.67, P=1.2×10^−30^, **Figure S7A**). This correlation was stronger in tumor (R=0.69, P=1.2×10^−13^) than normal samples (R=0.34, P=0.021). We hypothesized that average Tph usage would correlate with plasma cell abundance in tumors. To test this, we re-analyzed a published pan-cancer dataset containing other cell-types besides T-cells from 148 primary tumors, 53 matched adjacent tissues, and 25 healthy donor samples^62^. Tph usage and *CXCL13* expression remained correlated in this dataset (R=0.67, P=1.2×10^−30^, **Figure S7B**). Average Tph, Tfh-1, and Tfh-2 cGEP usage were significantly correlated with plasma cell percentage within the tumors (Spearman ρ=0.23, 0.34, 0.28, respectively, P<1×10^−2^, **Figure S7C**). In a multivariate regression across all samples, Tfh-1 and Tph usage were independently associated with plasma cell abundance (P=0.042, P=0.051 respectively). Subsetting to non-tumor samples, Tfh-1 and Tfh-2 remained statistically significant (P=0.017, P=0.027, respectively), but Tph was no longer significant (P=0.351). These findings suggest that Tph cells are functional within tumors and are associated with increased abundance of plasma cells.

## Discussion

Here, we introduced CellAnnoTator (abbreviated *CAT) for annotating scRNA-Seq data with predefined GEPs. *CAT exploits the observation that functionally informative GEPs learned by cNMF are reproducible across different datasets and contexts (**Figure 2**). This enables GEPs identified across multiple reference datasets to aid in interpreting new datasets. We demonstrated *CAT with a GEP catalog derived from T-cells across diverse tissues and diseases, yielding T-Cell AnnoTator (TCAT). We meta-analyzed a range of reference datasets, obtaining the most comprehensive T-cell GEP catalog to date, including 16 subset-associated, five technical artifact, and 25 functional programs.

TCAT demonstrated key advantages over clustering of T-cells. First, it simultaneously annotated functional and subset GEPs within the same cells, disentangling signals that clustering conflated (**Figure 4**). Second, TCAT out-performed RNA-based clustering for annotation of T-cell subsets without requiring manual curation of the cluster labels (**Figure 3**). Third, TCAT cGEP activity could be assessed across diverse disease states (**Figure 7**). TCAT also improved upon prior matrix factorizations of T-cells by yielding a more comprehensive catalog of T-cell GEPs. It was faster than running *de novo* matrix factorization, avoided the need to manually re-label GEPs, and increased accuracy for smaller datasets (**Figure 1C-F**).

TCAT explained why traditional T-cell subsets have been challenging to identify in scRNA-Seq. T-cell transcriptional clusters were heavily influenced by many non-subset GEPs, including technical artifacts, cell cycle, interferon response, and cytotoxicity (**Figure 4**). TCAT overcame this by annotating subset-associated cGEPs in parallel with functional cGEPs. In addition, TCAT revealed how cGEPs can be expressed in different contexts. For example, the cytotoxic cGEP was expressed in multiple subsets, and polarization cGEPs were expressed in both CD4 and CD8 T-cells (**Figure 4E, S4**). There has recently been increased recognition of polarized CD8 populations such as Tc2 which can secrete cytokines typically associated with Th2-polarized CD4 memory T-cells^40^. TCAT helped reveal these overlooked populations in scRNA-Seq data.

TCAT also highlighted the growing recognition of T peripheral helper (Tph) cells in disease. The Tph cGEP was significantly associated with Rheumatoid Arthritis (RA), Covid-19, and Cancer (**Figure 6**). While the association with RA was expected since Tph cells were discovered there, and recent data has identified Tph cells in Covid-19^39^, the association with cancer is less well established^63^. Tph usage was associated with expression of *CXCL13* and plasma cell abundance in tumors, suggesting Tph cells may drive lymphoid aggregation.

We also demonstrated that many cGEPs were induced following a TCR-dependent activation stimulus using the novel AIM-Seq assay (**Figure 5**). AIM-Seq produces TCR and CITE-Seq profiles for T-cells that are labeled based on their response to activation-induced marker assays. This identified 24 cGEPs associated with TCR-dependent activation, including 11 that may reflect context-dependent activation responses such as Th17-activated in Th17-polarized cells and CTLA4/CD38 in Tregs. Many of the AIM-associated GEPs were strongly associated with Covid-19, rheumatoid arthritis, and cancer, consistent with the importance of TCR-dependent activation in these diseases (**Figure 6**).

We aggregated several AIM-associated cGEPs into an antigen-specific activation (ASA) score to compare activation rates across diseases and cell subsets. This revealed impressive variability in the percentage of activated and exhausted CD4 and CD8 T-cells within and between different tumor types (**Figure 7**). In all tumor types, many T-cells lacked activation or exhaustion signatures and were labeled as bystanders. Bystanders were enriched for naive and unconventional T-cell subsets, whereas activated cells were enriched for Treg, Tph, and resident memory subsets. This approach shows how TCAT can aid in characterizing activation and exhaustion *in vivo*.

We highlight some current limitations of TCAT. First, TCAT’s output can be non-sparse, leading to non-zero usage of cGEPs contributing little biological function. This necessitates the use of thresholds balancing sensitivity and specificity to decide if a cGEP is active in a cell. For example, annotating TCR-activation or polarization currently relies on score thresholds. This limitation can be mitigated by algorithmic improvements that increase TCAT’s sparsity. Second, several cGEPs lack a clear interpretation, or may be redundant with other cGEPs in the catalog. For example, three cGEPs labeled IEG1-IEG3 are strongly enriched for immediate early genes. We used reproducibility of spectra across multiple datasets to enrich for biologically meaningful GEPs. As more datasets get incorporated, we anticipate increasing robustness of the catalog. Furthermore, new experimental perturbation datasets can facilitate linkage of cGEPs with upstream regulators to aid in interpretation.

We demonstrated application of *CAT to T-cells, but it is equally applicable to other cell types or tissues. We make the *CAT software publicly available and have created a repository to host cGEP catalogs, enabling easy application to new datasets. Furthermore, users studying other tissues and cell-types can contribute their own catalogs to the repository. We envision this as a resource akin to the molecular signatures database (MSigDB)^64,65^, but hosting GEPs for annotation of scRNA-Seq data rather than gene-sets for enrichment testing. We hope it will aid in comprehensive identification of GEPs underlying cell behavior across tissues and diseases.

## Methods

### Materials and reagents

| Reagent or Resource | Source | Identifier |
| --- | --- | --- |
| XVIVO15 culture media | Lonza | Catalog #: 02-060Q |
| RPMI 1640 Medium | ThermoFisher | Catalog #: 11875093 |
| Benzonase Nuclease | Sigma Aldrich | CAS #: 9025-65-4 |
| Anti-CD28 antibody | Biolegend | Catalog #: 302933<br>RRID: AB_11150591 |
| Anti-CD49d antibody | Biolegend | Catalog #: 304339<br>RRID: AB_2810443 |
| Human TruStain FcX™<br>(Fc Receptor Blocking<br>Solution) | Biolegend | Catalog #: 422302<br>RRID: AB_2818986 |
| Zombie Yellow™<br>Fixable Viability Kit | Biolegend | Catalog #: 423104 |
| TotalSeq™-C Human<br>Universal Cocktail, V1.0 | Biolegend | Catalog #: 399905 |
| Human TOTAL-SeqC<br>Repertoire (5') Hashing<br>Antibodies | BioLegend | Catalog #: 394661, 394663, 394665 |
| Anti-CD3-BV421 (SK7) | Biolegend | Catalog #: 344833<br>RRID: AB_2565674 |
| Anti-CD134-PE<br>(Ber-ACT35) | Biolegend | Catalog #: 350003<br>RRID: AB_10641708 |
| Anti-CD274-BV785<br>(29E.2A3) | Biolegend | Catalog #: 329735<br>RRID: AB_2629581 |
| Anti-CD137-APC<br>(4-B4-1) | Biolegend | Catalog #: 309809<br>RRID: AB_830671 |
| Anti-CD4-FITC<br>(RPA-T4) | Biolegend | Catalog #: 300505<br>RRID: AB_314073 |
| Chromium Next GEM Single Cell 5' Kit v2, 16 rxns | 10X | Catalog #:1000263 |
| Dual Index Kit TN Set A, 96 rxn | 10X | Catalog #: 1000250 |
| Chromium Next GEM Chip K Single Cell Kit, 48 rxns | 10X | Catalog #: 1000286 |
| Chromium Single Cell Human TCR Amplification Kit, 16 rxns | 10X | Catalog #: 1000252 |
| Library Construction Kit, 16 rxns | 10X | Catalog #: 1000190 |
| 5' Feature Barcode Kit, 16 rxns | 10X | Catalog #: 1000256 |

### CellAnnoTator (*CAT) Algorithm

Whereas cNMF learns both GEPs and their usage in cells, *CAT has the simpler problem of fitting the usage for a fixed set of GEPs. Specifically cNMF runs NMF multiple times, each time solving the following optimization:

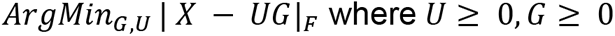

where *X* is a NxH matrix of N cells by the top H overdispersed genes, *U* is a learned NxK matrix of the usages of K GEPs in each cell, and *G* is a learned KxH matrix where each row encodes the relative contribution of each highly variable gene in a GEP. H is usually a parameter set to ∼2000 overdispersed genes. | |_F_ denotes the Frobenius norm. *X* includes variance-normalized overdispersed genes to ensure biologically informative genes are included and contribute similar amounts of information even when they may be expressed on different scales. For cNMF, the optimization is solved multiple times and the resulting *G* matrices are concatenated, filtered, and clustered to determine a final average estimate of *G*. Ultimately cNMF refits the GEP spectra into two separate representations, one reflecting the average expression of the GEP and units of transcripts per million *G*^*tpm*^ and on in Z-scored units used to define marker genes *G*^*scores*^ (see Kotliar et, al., 2019^14^ for details).

Analogously, *CAT takes a fixed catalog of GEPs as input, denoted as *G*^∗^, and a new query dataset *X*^*query*^ and solves the optimization:

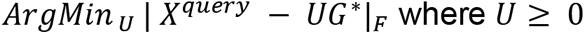

The columns of *X*^*query*^ and *G*^∗^correspond to a pre-specified set of overdispersed genes. Analogous to cNMF, we use gene-wise standard-deviation-normalized counts for *X*^*query*;^. See below for how *G*^∗^ is calculated for T-CellAnnoTator. We solve for *U* with non-negative least squares using the NMF package in scikit-learn version 1.1.3^66^ with *G*^∗^ fixed. We use the Frobenius error, the multiplicative update (“mu”) solver, tolerance of 1×10^−4^, and max iterations of 1000. We then row-normalize the *U* matrix so that each cell’s aggregate usage across all K GEPs sums to 1.

### Dataset pre-processing and batch-effect correction

To generate the input matrix for cNMF for each dataset, we first filtered genes detected in fewer than 10 cells and cells with fewer than 500 unique molecular identifiers (UMIs). We also excluded antibody-derived tags (ADTs) and genes containing a period in their gene name. We subsequently subsetted the data to the top 2000 most overdispersed genes, identified by the “seurat_v3” algorithm as implemented in Scanpy^35^. Next, we scaled each gene to unit variance. To avoid outliers with excessively high values, we calculated the 99.99th percentile value across all cells and genes and set this as a ceiling. We denote this matrix as *X*^*raw*^.

We used an adapted version of harmonypy to correct batch effect and other technical variables from *X*^*raw*^ prior to cNMF^21^. For this, we computed Harmony’s maximum diversity clustering matrix from principal components calculated from a normalized version of *X* which we label *X*^*norm*^. Specifically, to compute *X*^*norm*^, we started from the same initial gene list described above but first normalized the rows of the matrix so that each cell’s counts sum to 10,000 (TP10K normalization). We then subsetted to the top 2000 overdispersed genes, and scaled each column (gene) to unit variance, resulting in *X*^*norm*^. We then performed principal component analysis (PCA) on *X*^*norm*^ and supplied those principal components to the run_harmony function of harmonypy. We then used the mixture of experts model correction, implemented in harmonypy with the computed maximum diversity clustering matrix, but instead of correcting the PCs using this model, as standard Harmony does, we corrected *X*^*raw*^. This creates a small amount of variability around 0 for the smallest values in *X*^*raw*^. We therefore set a floor of 0, resulting in the corrected matrix *X*^*c*^ used as the count matrix for cNMF.

### Consensus non-negative matrix factorization (cNMF)

We ran cNMF on the batch-corrected *X*^*c*^ matrix which only includes the top 2000 overdispersed RNA genes. Spectra for the resulting GEPs were then refit by cNMF including all genes that passed the initial set of filters including ADTs. Specifically, RNA counts were normalized to sum to 10,000, and ADT counts were separately normalized to sum to 10,000 and the combined matrix was passed as the –tpm argument for cNMF. Thus the GEP spectra output by cNMF incorporate ADTs and genes not included in the 2000 overdispersed genes.

cNMF was run for each dataset with the number of components (K) varying between 15 and 55 and with 20 iterations. The final number of NMF components used for each dataset, K*, was chosen by visualizing the trade-off between reconstruction error and stability for these runs (**Supplementary item 1**). Once K* was selected, we ran cNMF a final time with only this value for K and with 200 iterations to generate the final GEP spectra estimates.

### Constructing a catalog of consensus GEPs (cGEPs)

Next, we identified consensus GEP spectra – I.e. the average of correlated GEP spectra identified by cNMF in different datasets. Normalized input GEP vectors, denoted as g_i_, were computed by starting from the spectra_tpm output from cNMF, renormalizing each vector to sum to 10^6^, and then dividing each element by the standard deviation of the corresponding gene in the –tpm input to cNMF. Then, we created an undirected graph where the 267 GEPs identified across all reference datasets were represented as nodes g_1_ … g_267_. We drew edges, denoted as E_i,j_ connecting a pair of GEPs g_i_ and g_j_ if the following criteria were met:

1. g_i_ and g_j_ were from different datasets
2. R_ij_ > 0.5 where R_ij_ denotes the Pearson correlation between g_i_ and g_j_. For computing R_ij_, g_i_ and g_j_ were subset to the union of the overdispersed genes for each dataset.
3. g_i_ was among the top seven most correlated GEPs with g_j_, and g_j_ was among the top seven most correlated GEPs with g_i_ with correlation defined as in 2.

Next, we initialized a set for each GEP: x_1_ = {g_1_} … x_267_ = {g_267_}. We then iterated through all edges E_i,j_ in the graph in order of decreasing R_ij_ and merged the sets x_i_ and x_j_ into a new set x_i,j_ = {g_i_, g_j_}. If either g_i_ or g_j_ were already members of a merged set from previous merges, we merged their containing sets only if at least two thirds of the GEP pairs in the resulting consensus set were connected by edges. For example, if there is an edge E_4,9_ and g_4_ is already merged into a set {g_1_, g_2_, g_4_}, then we only merged {g_1_, g_2_, g_4_} and {g_9_} if there were also edges E_1,9_ and E_2,9_. This resulted in 52 merged sets and 52 unmerged “singleton” sets. We filtered 49 of the 52 singletons and retained 3 that had a biological explanation for being identified in only one dataset.

Lastly, we subset each GEP to the union of overdispersed genes across all 7 reference datasets that were present in all dataset and obtained the final consensus GEPs by taking the element-wise average GEPs in each merged set. This matrix was used as the reference for TCAT. For marker gene analyses (e.g. **Figure 2B, D, Supplementary Item 2**), we element-wise averaged the Z-score representation of GEPs output by cNMF for GEPs in a consensus set.

### Simulation analysis

We adapted the scsim simulation framework described in the cNMF publication^14^ and based on Splatter^67^ into a new iteration, scsim2. Like with scsim, we distinguished between subset GEPs which are mutually exclusive and non-subset or “activity” GEPs which are not. For the original scsim framework, cells used one of multiple subset GEPs and potentially used a single activity GEP. We adapted scsim to allow cells to use anywhere from none to all of the activity GEPs in addition to their single subset GEP. We kept the Splatter parameters used in the cNMF publication to describe the distribution of gene expression data: mean_rate=7.68, mean_shape=0.34, libloc=7.64, libscale=0.78, expoutprob=0.00286, expoutloc=6.15, expoutscale=0.49, diffexpprob=.025, diffexpdownprob=.025, diffexploc=1.0, diffexpscale=1.0, bcv_dispersion=0.448, bcv_dof=22.087.

For figure 1, we simulated 10 subset GEPs and 10 activity GEPs based on 10,000 total genes. The extra-GEP reference included all 20, the missing-GEP reference included 6 of the subset GEPs and 6 of the non-subset GEPs, and the query dataset included 8 subset GEPs and 8 non-subset GEPs. Each dataset consisted of 9000 genes, randomly sampled from the 10,000. Each cell was randomly assigned a subset GEP with uniform probability (shown in the UMAP in figure 3B), and each cell randomly selected whether it expressed each activity GEP with probability of 0.3. The degree of usage of each activity GEP was sampled uniformly between 0.1 and 0.7. If the sum of the activity GEPs exceeded 0.8 for a cell, they were renormalized to sum to 0.8. Thus each cell’s usage of its subset GEP always exceeded 0.2. We simulated 100,000 cells each for the extra-GEP and missing GEP references. We simulated multiple query datasets containing 100, 500, 1000, 5000, 10,000, 20,000, 50,000, or 100,000 cells.

We subsequently ran cNMF using 1000 overdispersed genes, 20 iterations, local_neighborhood_size=0.3 and density_threshold=0.15. We used K=20, K=12, and K=16 for the extra-GEP reference, missing-GEP reference, and query datasets respectively. We then used *CAT to fit the usage of the reference GEPs on the query dataset. To evaluate the performance of *CAT and cNMF, we calculated the Pearson correlation of the inferred GEP usage with the simulated ground truth usage.

### Gene-set enrichment analysis

We used Fisher Exact Test in Python’s Scipy library to associate cGEPs with gene sets. For the T-cell polarization dataset^24^ we defined polarization gene sets as genes that had FDR-corrected P-value < 0.05 and fold change > 2 with the stimulation condition. We excluded genes with FDR-corrected P-value between 0.05 and 0.2 and fold-change>1, as many of these are up-regulated by the stimulation but just did not reach FDR significance. We also obtained literature gene sets corresponding to immediate early genes^28^ and gene ontologies^23,68^. We tested these literature gene-sets for enrichments with gene sets derived from the Z-score representation of cGEPs based on a score threshold of 0.015, which corresponded to the 99th percentile across all genes and cGEPs. We then tested for association using Fisher’s Exact Test as implemented in scipy.stats in Python.

### Manual subset gating analysis

We library-size normalized antibody derived tag (ADT) protein measurements to sum to 10^4^ (TP10K) and applied the centered log ratio (CLR) transformation. We then scaled each protein to unit variance, and truncated at 15 to remove excessively high outliers. Next, we performed principal component analysis (PCA) and ran batch correction using harmonypy with the same batch features as for cNMF. We then computed the K-nearest neighbor graph with K=5 neighbors, using the Harmony-corrected principal components. We then smoothed the normalized protein estimates using MAGIC^69^ using the K-nearest neighbor graph computed above and the diffusion operator powered to t=3.

We gated canonical T-cell subsets using the smoothed normalized ADTs. First, we gated gamma-delta (γδ) T-cells using expression of Vδ2 TCR. Then, we separated MAIT cells using expression of CD161 and TCR Vα 7.2. We then used CD4 and CD8 to separate CD4 (CD4+CD8-), CD8 (CD4-CD8+), double positive (DP) (CD4+CD8-), and double negative (DN) (CD4-CD8-) T-cells. We then subset to CD4 T-cells and gated regulatory T-cells (Tregs) using expression of CD25 and CD39. Of the remaining CD4 T-cells, we used CD62L and CD45RA to define CD4 Naive (CD62L+CD45RA+), CD4 Central Memory (CD62L+CD45RA-), CD4 Effector Memory (CD62L-CD45RA-), and CD4 TEMRA (CD62L-CD45RA+) populations. For the CD8 T-cells, we similarly used CD62L and CD45RA to define CD8 Naive (CD62L+CD45RA+), CD8 Central Memory (CD62L+CD45RA-), CD8 Effector Memory (CD62L-CD45RA-), and CD8 TEMRA (CD62L-CD45RA+) populations.

### T-cell subset classification benchmarking analyses

We used T-cell subsets defined by manual gating of ADTs in the Flu-Vaccine dataset as ground truth for prediction. For single cGEP prediction, we ran TCAT to predict cGEP usage, and identified the cGEP that best predicted the lineage based on area under the curve (AUC).

We also used all of the cGEP simultaneously to perform simultaneous multi-label prediction. We scaled the normalized usages for all cGEPs to zero mean and unit variance. Using COMBAT as a training dataset, we trained a multinomial logistic regression using scikit-learn^66^ version 1.0.2 with lbfgs solver to predict gated subset from usages. Model weights were adjusted by the inverse of subset size using class_weight=“balanced”, allowing subsets with different cell counts to contribute to the model equally. We excluded CD4 TEMRA, double negative, and double positive subsets from this analysis due to low cell counts in both the training and testing datasets. We evaluated this model in the independent Flu-Vaccine query dataset.

Analogous comparisons were made using GEPs from Yasumizu et. al, 2024 fit to the data using the NMFproject software^13^. We also obtained gene sets derived from NMF analyses of T-cell in a pan-cancer dataset^16^. To assess the ability of these gene sets to predict gated subsets, we used the score_genes function in Scanpy^35^ on data normalized following the standard pipeline (library size normalizing to TP10K, log transformation, scaling each each gene to unit variance). We then assigning each subset to the gene set that yielded the maximal AUC.

To evaluate clustering, we first normalized the data as above, and subset to highly variable genes using the highly_variable_genes function in Scanpy with default parameters. We then ran principal component analysis (PCA) and Harmony batch correction of the PCs^21^. We then computed the K nearest neighbor graph using 31 harmony-corrected PCs and 30 nearest neighbors. We then performed Leiden clustering^70^ with resolution parameters ranging from 0.25 to 2.25 increasing by 0.25. For each clustering resolution, we performed a greedy search to assign clusters to manually gated subsets based on maximization of the balanced accuracy (I.e. the average recall across all subsets). In each iteration, we considered all unassigned clusters and possible gated subset assignments, and selected the cluster and assignment that most increased the overall balanced accuracy. When no remaining cluster assignments would increase the balanced accuracy, we assigned the cluster to a subset that least decreased the balanced accuracy. We continued this process until each cluster was assigned to a subset.

### Activation Induced Marker assay followed by scRNA-Seq (AIM-Seq)

PBMCs were quickly thawed and placed in pre-warmed xVIVO15 cell culture medium (Lonza) supplemented with 5% heat-inactivated FBS. To reduce cell clumping, PBMCs were incubated in xVIVO15 containing 50 U/mL of benzonase nuclease (Sigma-Aldrich) for 15 minutes at 37 degrees and filtered using a 70 µm cell strainer. Washed and nuclease treated cells were seeded in a 96 well cell culture plate at a concentration of 2.5 x 10^6^/mL. Peptide stimulations were performed using the CEFX Ultra SuperStim Pool (JPT Peptide Technologies, Product Code: PM-CEFX-1) at a final concentration of 1.25 µg/mL per peptide for 22 hours at 37 degrees and 5% CO_2_. Recombinant anti-CD28 and anti-CD49d antibodies (BioLegend) were added at a final concentration of 5 µg/mL and 0.625 µg/mL, respectively, to provide co-stimulation for peptide reactive T-cells. Separately mock-stimulated cells were treated with anti-CD28 and anti-CD49d antibodies at the same concentration.

Peptide responsive T-cells were detected by the expression of the surface activation markers PD-L1, OX40, and CD137 via flow cytometry. Following the stimulation, peptide treated and mock-stimulated cells were washed in cell staining buffer (PBS + 2mM EDTA + 2% FBS) to end the stimulation. Fc receptor blocking was performed using a 1:50 dilution of Human TruStain FcX (Biolegend) in cell staining buffer for 10 minutes at 4 degrees. Cell viability staining was performed using a 1:500 dilution of Zombie Yellow Fixable Viability Dye (BioLegend) prepared in PBS for 30 minutes at 4 degrees. Surface staining was performed using 1:100 dilutions of BV421 conjugated anti-CD3, FITC conjugated anti-CD4, BV786 conjugated anti-PD-L1, PE conjugated anti-OX40, and APC conjugated anti-CD137 (BioLegend) for 25 minutes at 4 degrees in cell staining buffer. Following cell staining, antigen reactive and non-reactive T-cells were identified using a BD FACSAria II cell sorter and collected in cRPMI medium (100 U/mL penicillin-streptomycin + 2 mM L-glutamine + 10 mM HEPES + 0.1 mM non-essential amino acids + 1 mM sodium pyruvate + .05 mM 2-Mercaptoethanol) supplemented with 20% FBS. Sorted T-cell populations were then labeled with 75 uL of TotalSeq oligo conjugated hashing antibody mix, incubated for 30 minutes at 4 degrees with gentle mixing after 15 minutes, and pooled in equal quantities. Staining with the TotalSeq-C Human Universal Cocktail (BioLegend) was then performed according to the manufacturer’s instructions. The cells were then resuspended in PBS supplemented with .04% FBS at a final concentration of 500 cells/µL and submitted for single-cell profiling on the Chromium Next GEM instrument. Library preparation was completed for the hashtag oligos, single-cell rna-seq, cite-seq, and TCR-repertoire sequencing following the manufacturer’s instructions.

We collected AIM-Seq data from two separate 10X runs. In the first experiment, PBMCs from three donors were processed independently as described above and were pooled together after fluorescence activated cell sorting (FACS). In the second run, PBMCs from four donors, two of which overlapped with the first run, were stimulated separately and pooled prior to FACS.

### Preprocessing the AIM-Seq dataset

The AIM-Seq data was processed using Cell Ranger version 6.1.1 with default parameters and alignment to hg38 reference genome. The donor of origin for each cell was determined using Demuxlet version 1.0 with doublet-prior of 0.1^71^. Cells with null or ambiguous demuxlet result, fewer than 10 counts of the hashtag oligos, or fewer than 50 total RNA counts were filtered. To account for staining differences between the hashtag oligos and different sequencing depths of the two 10X runs, the counts for each hashtag oligo in each 10X run were scaled to have the same median value. Next we added a pseudocount to the hashtag oligo counts and log10 transformed this data. Then we ran Gaussian Mixture models separately for each hashtag oligo with K=2 clusters. Each cell was assigned to a single condition if it was in the high cluster for one oligo and the low clusters for all others, a doublet if it was in the high cluster for more than one oligo, or an empty droplet if it was in the low cluster for all oligos. Empty droplets or doublets based on the hashtag oligo clustering were filtered, as were doublets based on demuxlet. Genes detected in fewer than 10 cells were filtered prior to running TCAT.

### cGEP associations with AIM-positivity, proliferation, and disease

To associate cGEPs with the AIM-Seq stimulus, we first ran TCAT to fit the usages of the cGEPs in the AIM-Seq dataset. We then computed the average usage of each cGEP in cells from each sort condition in each donor. We created two dummy variables, the first indicating whether a sample was treated with CEFX or Mock, and the second indicating whether a CEFX-treated sample was AIM-positive or not. We fit these two variables and an intercept to average cGEP usage in the sample. cGEPs associated with the CEFX-or-Mock dummy variable were labeled milieu-associated while cGEPs positively associated with the AIM-positive dummy were labeled AIM-associated.

To associate cGEPs with proliferation, we defined cells as proliferating or non-proliferating in each dataset by setting a threshold of 0.1 on the sum of the three cell cycle cGEPs, S-phase, late S-phase, and G2M-phase. We then computed the mean usage of each cGEP per sample separately in high cell-cycle (sum usage > 0.1) and low cell-cycle (sum usage < 0.1) cells. We filtered samples that did not have at least 10 high cell-cycle cells and 100 low cell-cycle cells. Then, for each cGEP, we performed a two-sample T-test paired by individual (ttest_rel in Scipy, default parameters) between average cGEP usage for high and low cell-cycle cells. We meta-analyzed P-values across datasets using Fisher’s Method (combine_pvalues in Scipy).

To associate cGEPs with sample-level disease phenotypes, we calculated the average usage of each cGEP in each sample for a given dataset. We then used ordinary least squares regression to find cGEPs with higher average usage in disease samples than controls, controlling for sample-level batch variables as covariates. For all datasets, disease status was modeled as a binary dummy variable, and an intercept was included. For UK-Covid, the processing site was included as dummy variable covariates. For COMBAT, sequencing pool, and processing institute were included as dummy variable covariates. For the Pan-cancer dataset, all cancer types were initially included in the analysis and dummy variable covariates were included for tissue of origin. In addition, sequencing technology was included as a dummy variables. When there were multiple tumor samples or matched normal samples from the same donor, we retained only the duplicate sample with the most cells prior to the regression.

For all association tests, we performed FDR-correction of the P-values using the Benjamini Hochberg method (fdrcorrection in Statsmodels with method=’indep’).

### Defining the antigen-specific activation (ASA) score

We used CD71+CD95+ surface protein co-expression in the COMBAT and Flu-Vaccine datasets as an *in vivo* correlate of TCR activation to help prioritize AIM-associated cGEPs for predicting TCR-activated cells. First we preprocessed the ADT surface proteins in these datasets as described in the manual subset gating section. We then subsetted cells by their manual gating-defined broad cell types (CD4 Conv, CD4 Treg, CD8 Conv, other) and gated CD71+CD95+ cells separately for each cell type as the response feature to be predicted by AIM-associated cGEPs.

We then performed forward stepwise selection, evaluating how well the summation of usages of different combinations of AIM-associated cGEPs would predict CD71+CD95+ gating. At each stage, the per-cell ASA score was computed as the sum of normalized usages of cGEPs in the predictive set. At each forward step, we determined which cGEP should be added to the predictive set based on which would most improve the average AUC across the Flu-Vaccine and COMBAT datasets. We used a reduction in AUC in both datasets as the stopping criterion for adding cGEPs. We considered all AIM-associated cGEPs identified in section 6 as candidates for this, excluding those known to have a broader function outside of T-cell activation (e.g. cytoskeleton, metallothionein, cell cycle) and those reflecting activation-associated T-cell subsets (Tph and Th17-Activated). We also excluded Exhaustion from the ASA score as it reflects a distinct inhibitory response to antigen-stimulation that users may wish to annotate separately.

### Code availability

The code for CellAnnotator (starCAT) is available at https://github.com/immunogenomics/starCAT. The analysis scripts used in this paper are available at https://github.com/immunogenomics/TCAT_analysis.

## Acknowledgements

This work was supported by the following NIH grants: P01AI148102, UCAR081023, U01HG012009, R01AR063759, and R56HG013083. P.C.S. is supported by the Howard Hughes Medical Institute. K.W. is supported by NHGRI T32HG002295 and NIAMS T32AR007530.

## Author contribution

Conceptualization (D.K., M.C., S.R.), Software (D.K., M.C.), Formal Analysis (D.K., M.C.), Methodology (D.K., M.C., R.A., S.R.), Writing (D.K., M.C., K.W., A.N., Y.B., Y.Z., P.C.S., D.A.R., S.R.), Funding acquisition (S.R.)

## Tables / Legends

**Table S1. cGEP Summary.** Summary of cGEPs including their full name, abbreviated name, assigned class, top 3 most strongly associated genes, and which datasets it was derived from.

**Table S2. Marker genes.** Top 200 marker genes associated with each cGEP, colored by their strength of association with the cGEP, based on the average gene score.

**Table S3. Gene-set enrichment.** The “GO_Enrichment” tab includes the top 10 associated gene sets for each cGEP including the GEP name, gene-set name, fisher exact test odds ratio, and P-value. The subsequent tabs include the same information but for enrichment tests for gene sets defined from a dataset that polarized T-cells for either 16 hours (16h) or 5 days (5d) starting from either naive (TN) or memory T-cells (TM)^24^. The tab name indicates the stimulation conditions.

**Table S4. Correlation with cell quality features.** Each tab includes the Pearson correlation of each cGEP’s usage (rows) with different per-cell quality features (tab names) for each dataset (columns). MitoFrac denotes the % of unique molecular identifiers from MT-genes. RNA_Detected denotes the number of unique genes detected per cell. RNA_Count denotes the number of unique molecular identifiers per cell. PCFrac denotes the percentage of unique molecular identifiers that are assigned to a protein coding gene in Gencode version 44.

**Table S5. AIM-Seq association.** Provides regression coefficients and P-values for the association between cGEP usage and binary variables reflecting CEFX vs. mock stimulation or AIM-positive vs. AIM-negative. Coef. represents the regression coefficient, P represents the P-value, and Q represents the FDR-corrected P-value.

**Table S6. Association with proliferation.** T-statistics, P-values, and log_2_ odds ratios for the paired T-test of proliferating and non-proliferating T-cells in each dataset (tabs). For the meta-analysis across datasets it provides the Fisher’s method combined P-value and the average log_2_ odds ratio.

**Table S7. Association with disease.** ordinary least squares regression coefficients (Beta), P-values (P), FDR-corrected Q-values (Q), and average fold changes (FC) for phenotype associations shown in **Figure 7**. Each tab represents a different phenotype.

## Supplemental Figures / Legends

**Figure S1.**
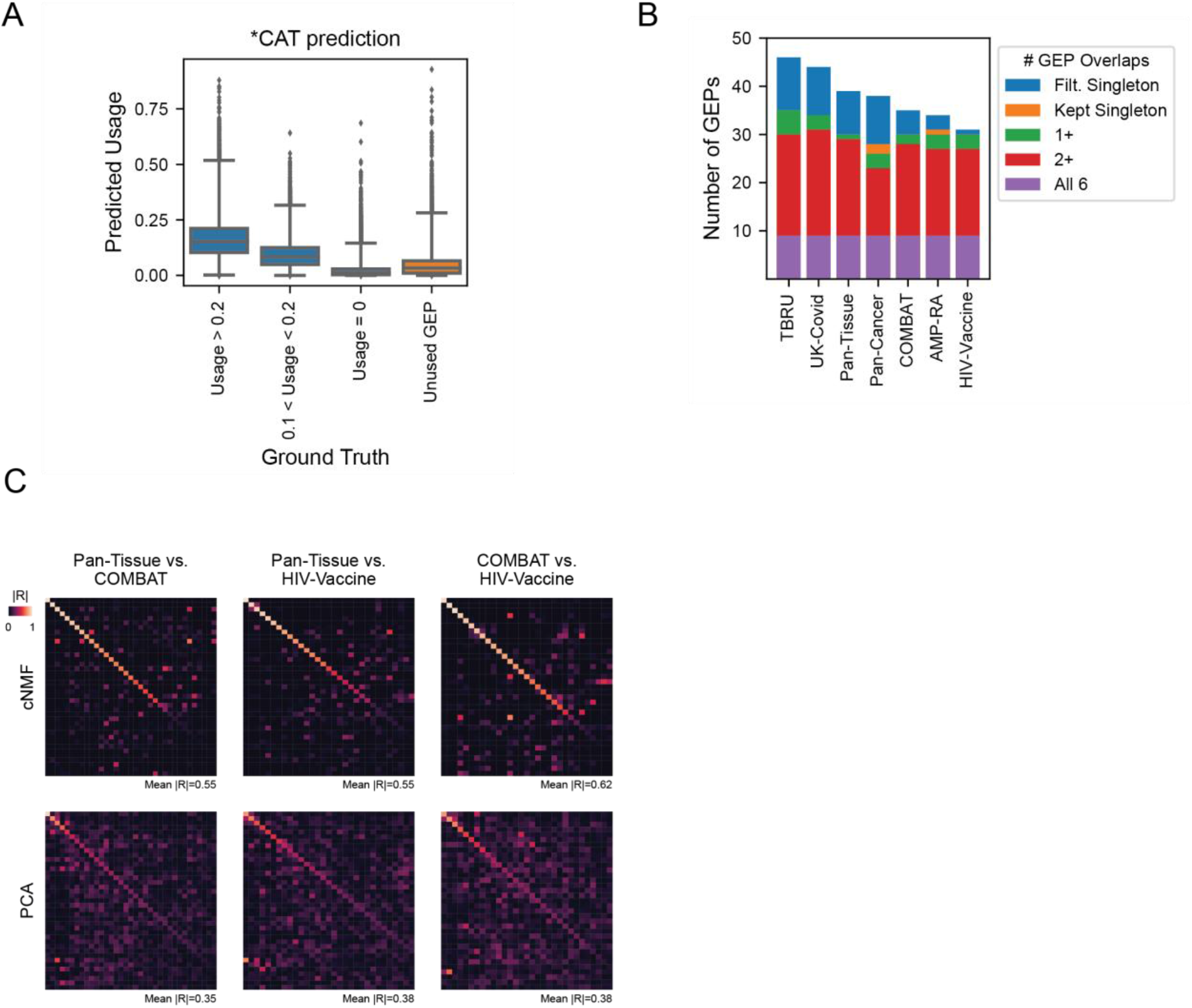
Characterizing *CAT. (A) *CAT predicted GEP usage for cells that use a GEPs with ground-truth usage>0.2, 0.1-0.2, or 0. Also shows the predicted usage for GEPs present in the reference data that are not present in the query (labeled unused GEP). (B) Number of GEPs identified in each dataset. The color indicates whether each GEP clustered with one or more GEPs from another dataset as part of a consensus GEP (purple, red, or green), did not cluster with a GEP from another dataset but was kept in the catalog as a dataset-specific GEP (orange), or did not cluster with a GEP from another dataset and was filtered (blue). (C) Absolute value of Pearson correlation of spectra learned by cNMF (top) or PCA (bottom) between different pairs of datasets. PCs are learned on the same matrices of batch-corrected matrices used for cNMF. Mean correlation refers to the mean value along the matrix diagonal, which corresponds to pairs of components with highest correlation across the two datasets.

**Figure S2.**
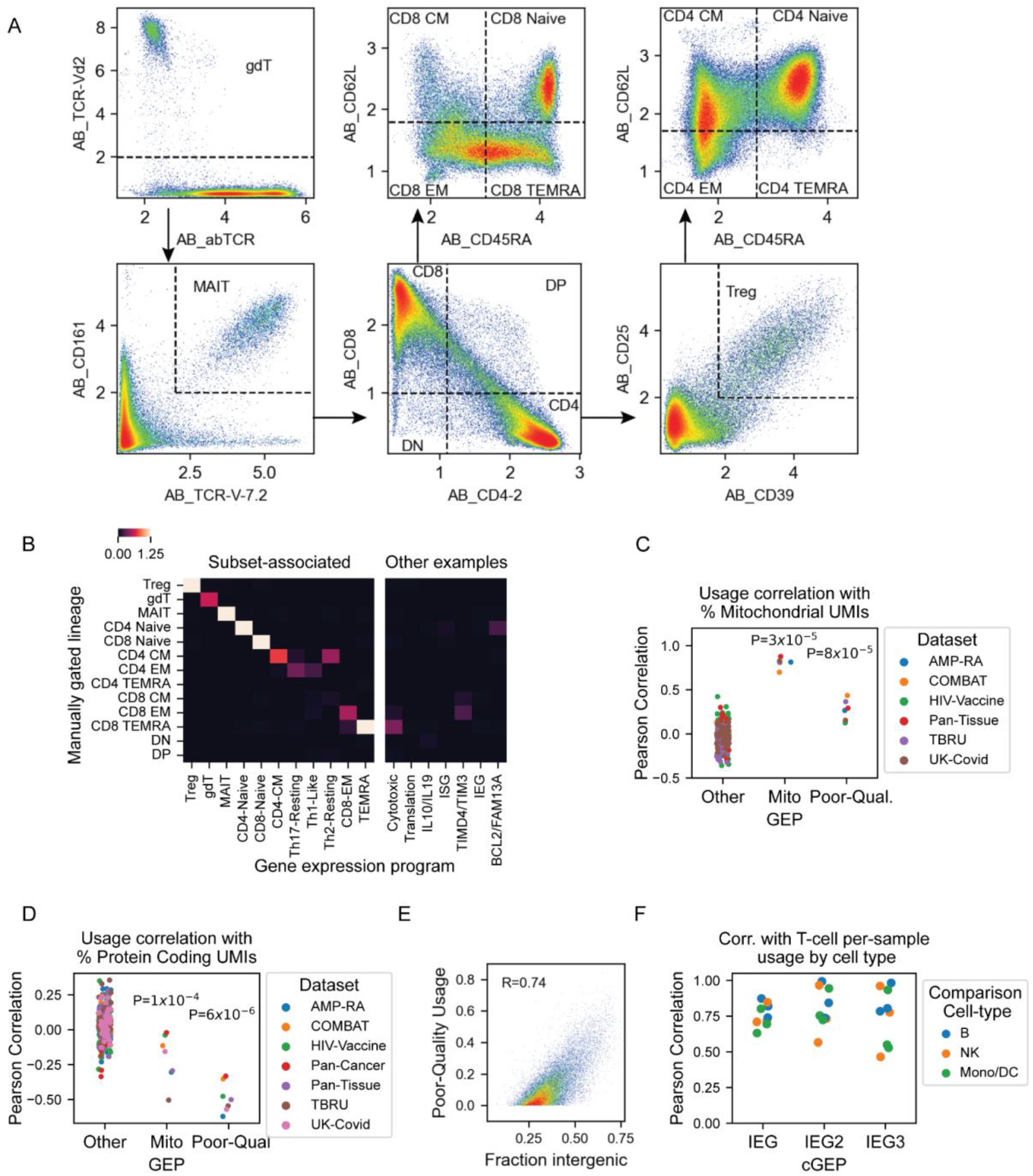
Annotating cGEPs. (A) Manual gating of COMBAT dataset using smoothed surface protein antibody-derived tag (ADTs). (B) Multivariate logistic regression coefficients of cGEPs (columns) against manually gated populations (rows). For visualization, the minimum value is thresholded to 0 and the maximum is threshold to 1.25. Seven selected non-subset cGEPs are shown on the right as examples. (C) Pearson correlation of cGEPs with percentage of mitochondrial transcript per cell, for each dataset. All cGEPs excluding Mito and Poor-Quality are included in the “Other” column. P-values are from a Ranksum test of the selected cGEP against the Other cGEPs. (D) Same as (C) but showing correlation with the percentage of UMIs assigned to protein coding genes. (E) Scatter plot of the proportion of UMIs mapping to intergenic regions in the genome against Poor-Quality cGEP usage for cells in the AMP-RA dataset. (F) Correlation of per-sample average cGEP usage in T-cells with that in B-cells, NK-cells for the 3 immediate early gene cGEPs, in the COMBAT, UK-Covid, and HIV-Vaccine datasets.

**Figure S3.**
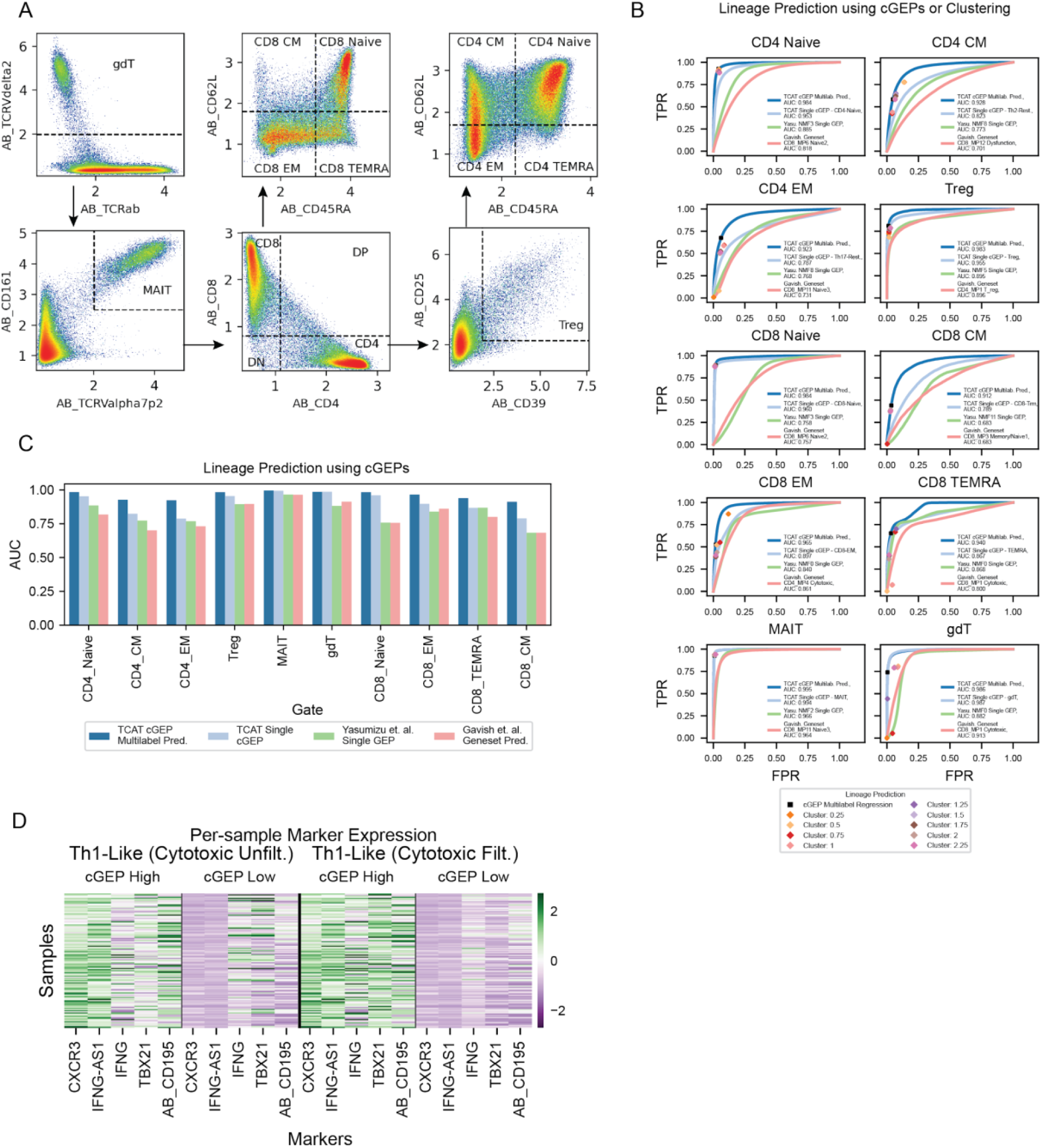
Benchmarking CellAnnoTator on simulated and real datasets. (A) Manual gating for the Flu-Vaccine dataset analogous to **Figure S2A**. (B) Receiver operator curves (ROCs) for prediction of manually gated subset based on a single most associated subset (dark blue), TCAT multilabel prediction (light blue), analogous predictions using the single most associated NMF component published in Yasumizu et al., 2024^13^, or using gene sets from NMF components in Gavish et al., 2023^16^. Individual points show accuracies of discrete predictions based on cGEP multilabel regression, or clustering with the leiden resolution specified in the legend. (C) Areas under the curve (AUC) from receiver operator curves in (B). (D) Heatmap of pseudobulk expression in Th1-Like-high and low cells, per sample. Cytotoxic-high cells are included (left) and filtered (right). Sample expression is normalized by library size and z-scored across rows, separately for the two filtering conditions.

**Figure S4.**
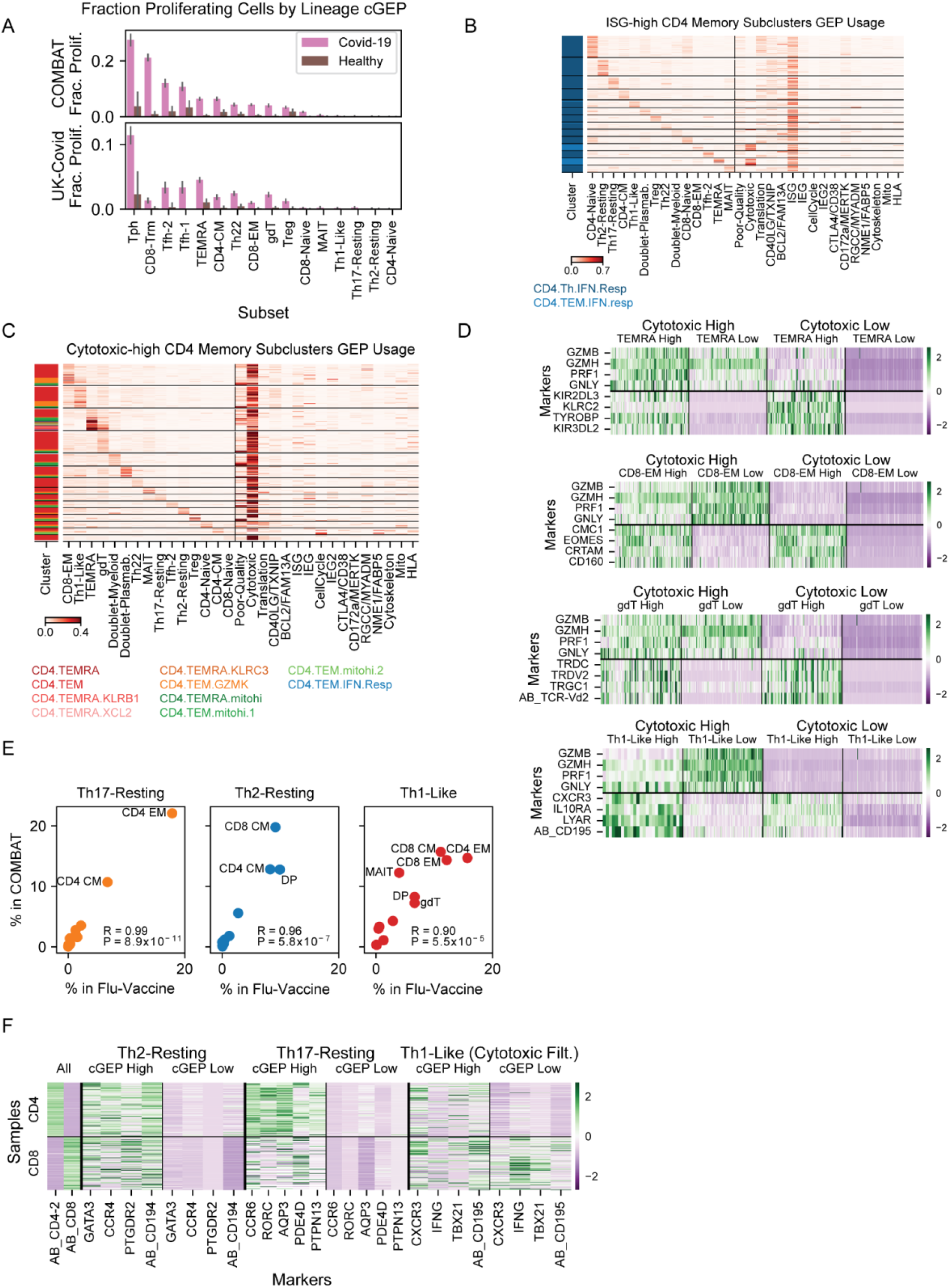
Comparing TCAT with COMBAT dataset clustering. (A) Fraction of proliferating cells (cell cycle usage>0.1) assigned to each subset based on the most highly used subset-associated GEPs, for cells from Covid-19 or healthy donors in the two Covid-19 datasets. Error bars represent 95% bootstrap confidence intervals. (B) Usage of selected cGEPs (columns) in cells (rows) grouped by maximum subset cGEP. Cells are drawn from subclusters with high usage of the ISG cGEP, indicated in the colorbar. (C) Same as (B) but only showing cells from subclusters with high cytotoxicity cGEP usage. (D) Heatmap of pseudobulk expression of marker genes in cytotoxic-high and low cells and subset cGEP high and low cells, per sample. Expression is normalized by library size and z-scored across rows. (E) Average fraction of polarized cells (usage>0.1) per gated subset, across samples, within COMBAT and Flu-Vaccine datasets. (F) Heatmap of pseudobulk expression of marker genes in polarization-high and low cells, separately for gated CD4 and CD8s T-cells, per sample. Sample expression is normalized by library size and z-scored across rows, for each polarization.

**Figure S5.**
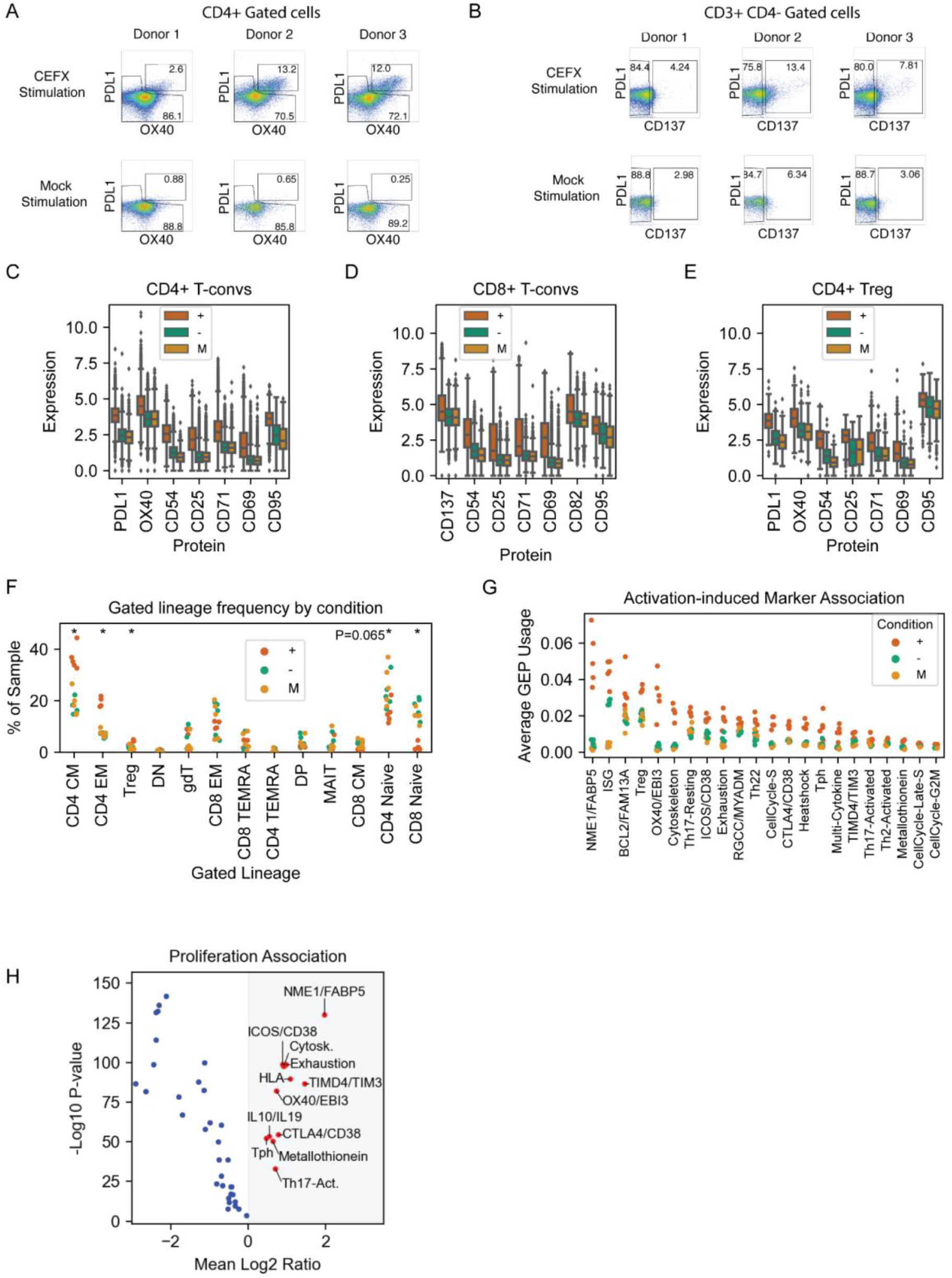
Identifying activation associated cGEPs with AIM-Seq. (A-B) Flow cytometry data of CD3+CD4+ and CD3+CD4-gated populations for 3 donor samples for CEFX and mock conditions. (C-E) Activation-induced marker (AIM) surface protein expression based on CITE-Seq for CD4+, CD8+, and Treg subsets, stratified by sort condition. Boxes represent interquartile range and whiskers represent 95% percentiles. (F) Percentage of each sample assigned to each subset based on manual gating, colored by stimulation condition. * indicates t-test P<.05 comparing + and U. (G) Average cGEP usage in each donor and condition, for AIM-associated cGEPs. (H) Paired t-test of pseudobulk cGEP usage in high and low cell cycle usage cells (threshold 0.1) from each sample. X-axis shows the mean Log_2_ ratio of average usages across datasets. Y-axis shows the –Log_10_ P-value. Statistically significant and positively associated cGEPs are indicated in red.

**Figure S6.**
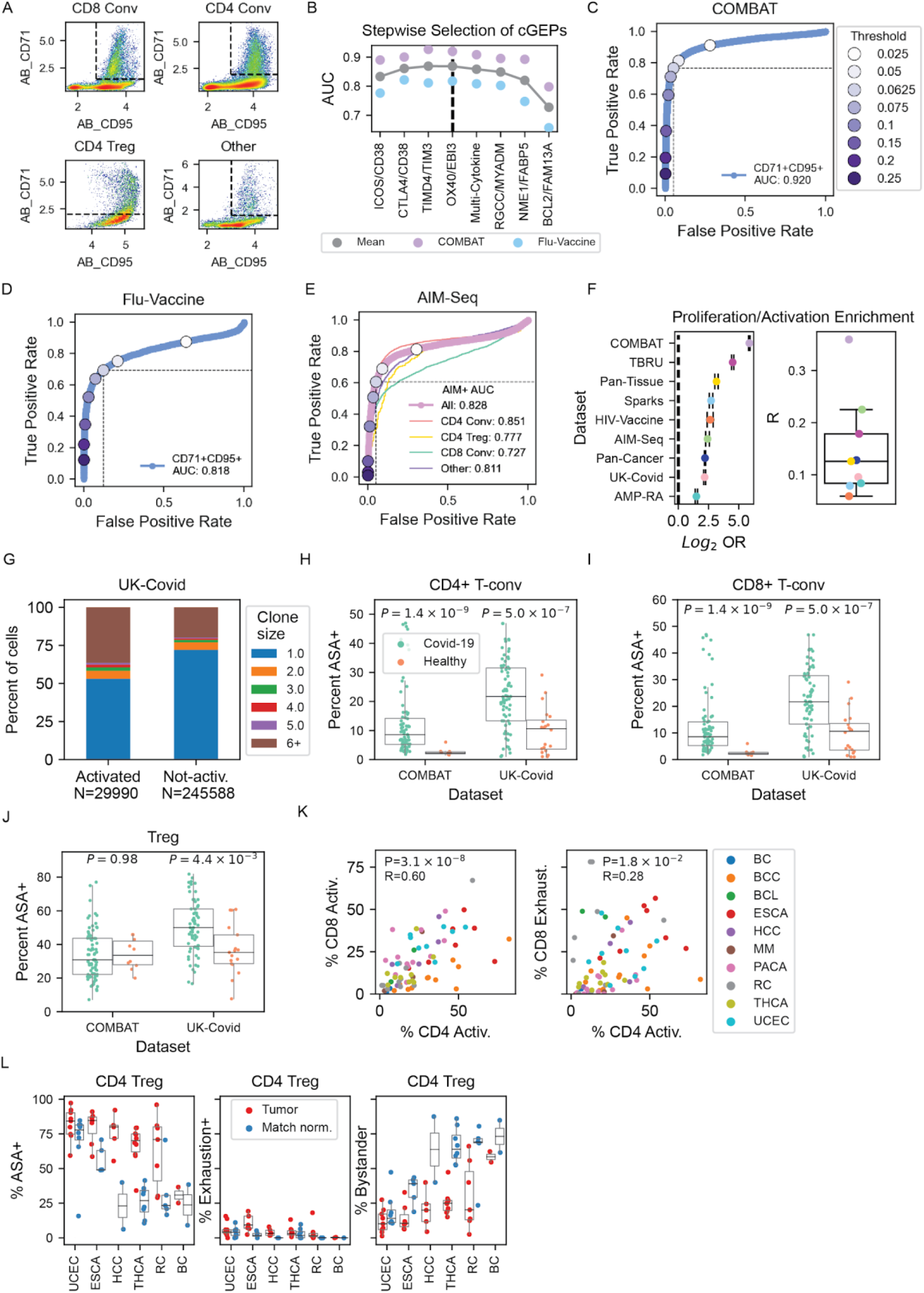
Annotating antigen-specific activation *in vivo*. (A) Definition of activation used for training the antigen-specific activation (ASA) score in the COMBAT dataset for manually gated subsets. (B) AUC estimates averaged for predicting CD71/CD95 co-expression based on summation of cGEPs sequentially added to the score from left to right. (C-D) Receiver operator curve (ROC) for ASA prediction of CD71/CD95-based activation labels, with various thresholds denoted as colored points. (E) ROC for ASA prediction of AIM-positivity in the AIM-Seq dataset. (F) Left – Odds ratio of enrichment between proliferation (aggregate cell cycle cGEP usage>0.1) and activation (ASA>0.065) for each dataset. Error bars denote 95% confidence intervals. Right – Pearson correlation between ASA and aggregate cell cycle cGEP usage with colors mapping to dataset. (G) Clonality in manually gated conventional CD4 and CD8 T-cells annotated as activated (ASA>0.065) or not activated (ASA<0.065). Clonality is defined as the number of cells in the same sample with an identical alpha and beta CDR3 amino acid sequence. (H-J) Percentage of activated CD4 convs, CD8 convs, and Tregs based on ASA>0.065 in Covid-19 and healthy control samples from COMBAT and UK-Covid datasets. (K) Percentage of activated conventional CD4 T-cells (ASA>0.065) versus percentage of activated or exhausted (exhaustion usage>0.065) conventional CD8 T-cells across tumor samples. (L) Percentage of activated, exhausted, or bystander (ASA + exhaustion usage<0.065) Tregs in tumors and match normal samples.

**Figure S7.**
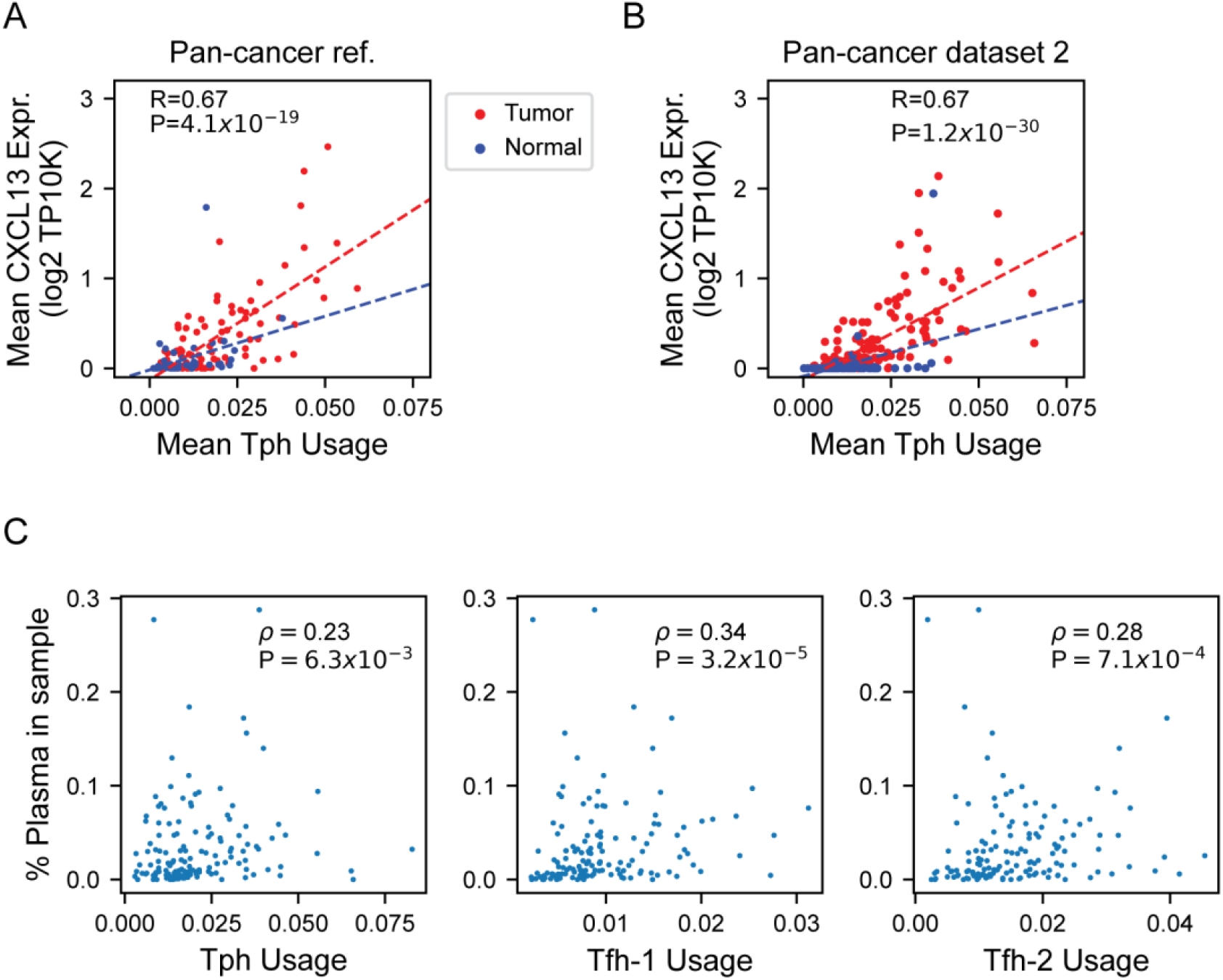
Identifying cGEPs associated with disease phenotypes. (A-B) Average usage of the T peripheral helper (Tph) cGEP compared to average *CXCL13* expression from T-cells within tumors and matched normal tissue samples in Pan-cancer reference and Luo et al., 2022^62^. Trend lines show the regression coefficients fit for tumors and normal samples separately (D-F) Percentage of cells annotated as plasma cells against the average Tph, Tfh-1, or Tfh-2 usage within T-cells from tumor samples in Luo et al., 2022^62^.

## Supplementary Item Figures / Legends

**Supplementary item 1.**
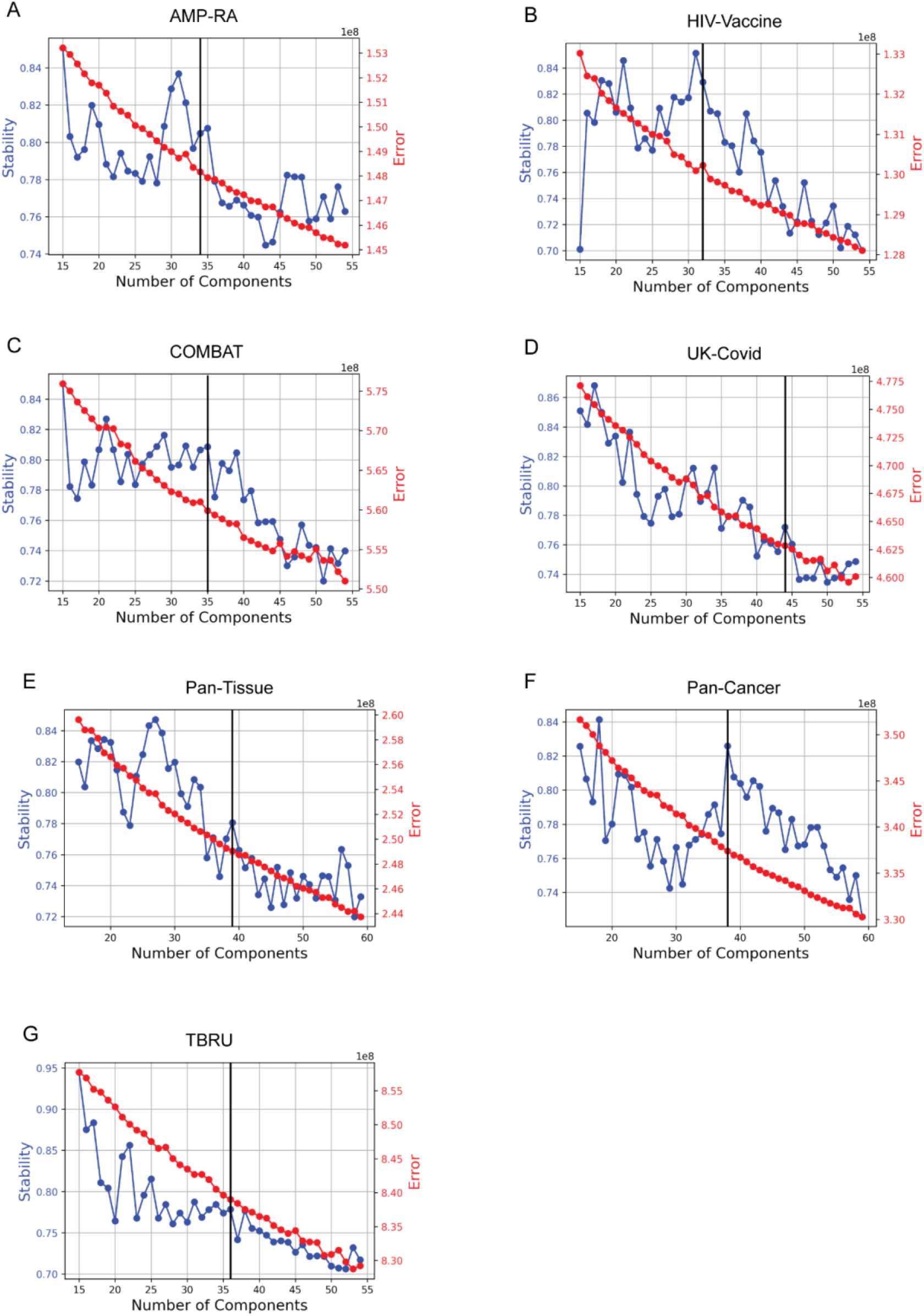
K selection plots for consensus NMF runs on reference datasets. Vertical line denotes the selected number of components.

**Supplementary item 2.**
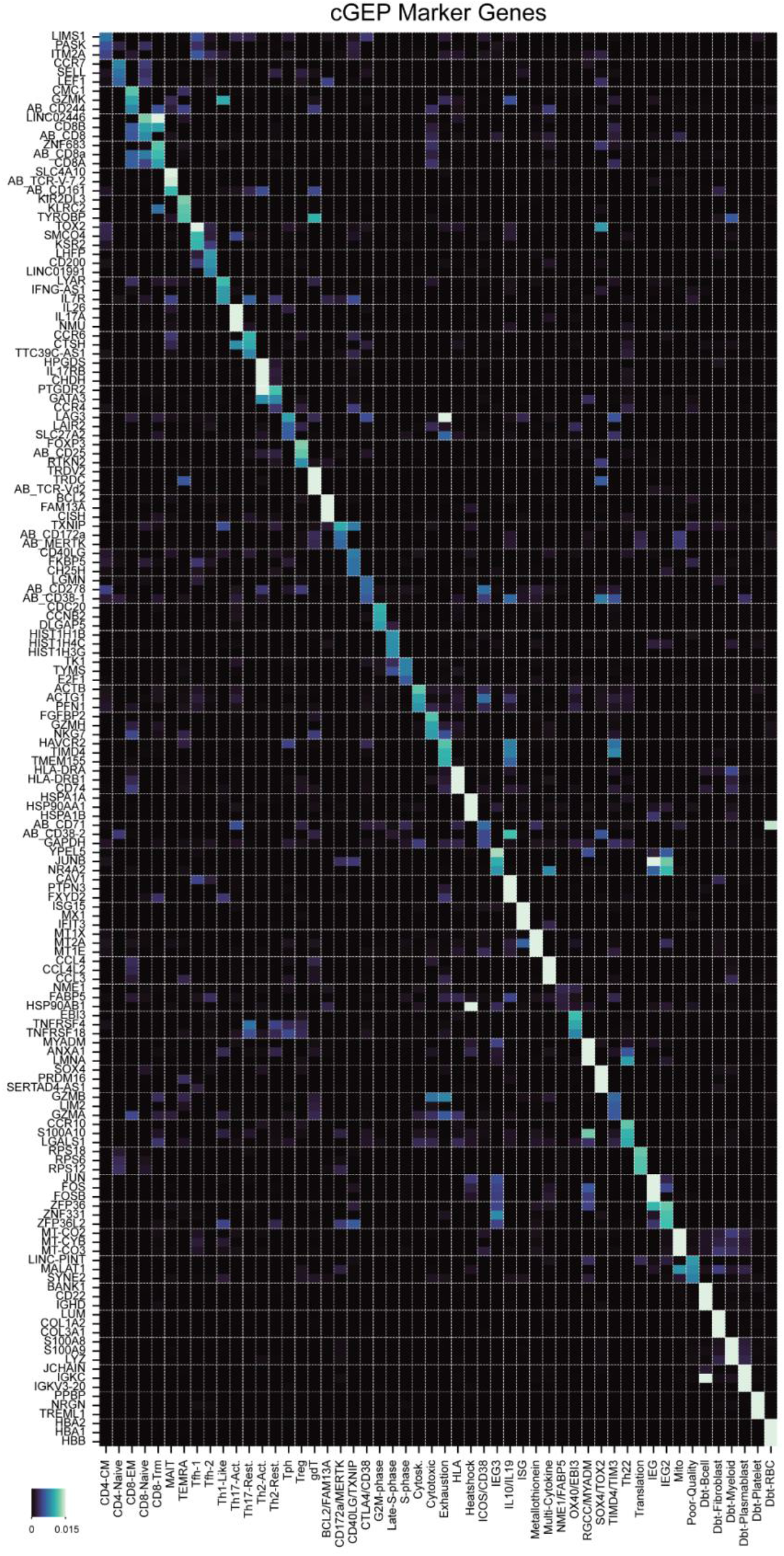
Example marker genes for all cGEPs. Color indicates average cNMF gene score units which denotes how much 1 additional count of usage of the cGEP would be expected to increase expression of the gene in Z-scored units.

**Supplementary item 3.**
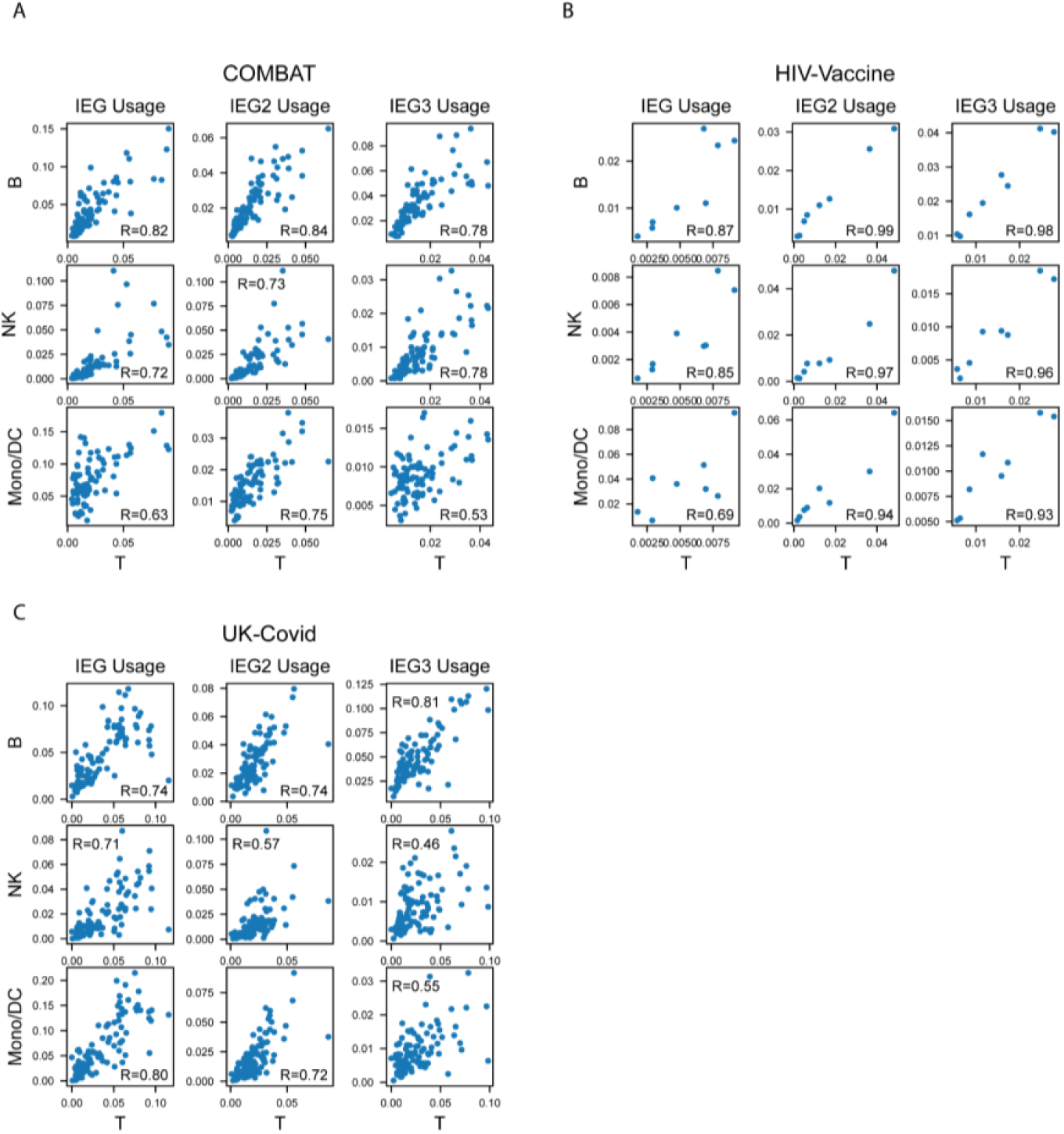
Immediate early gene usage across circulating blood cell types. Average per-sample usage of each IEG cGEP in T-cells versus monocytes and dendritic cells, NK cells, or B-cells, in the three reference PBMC datasets.

**Supplementary item 4.**
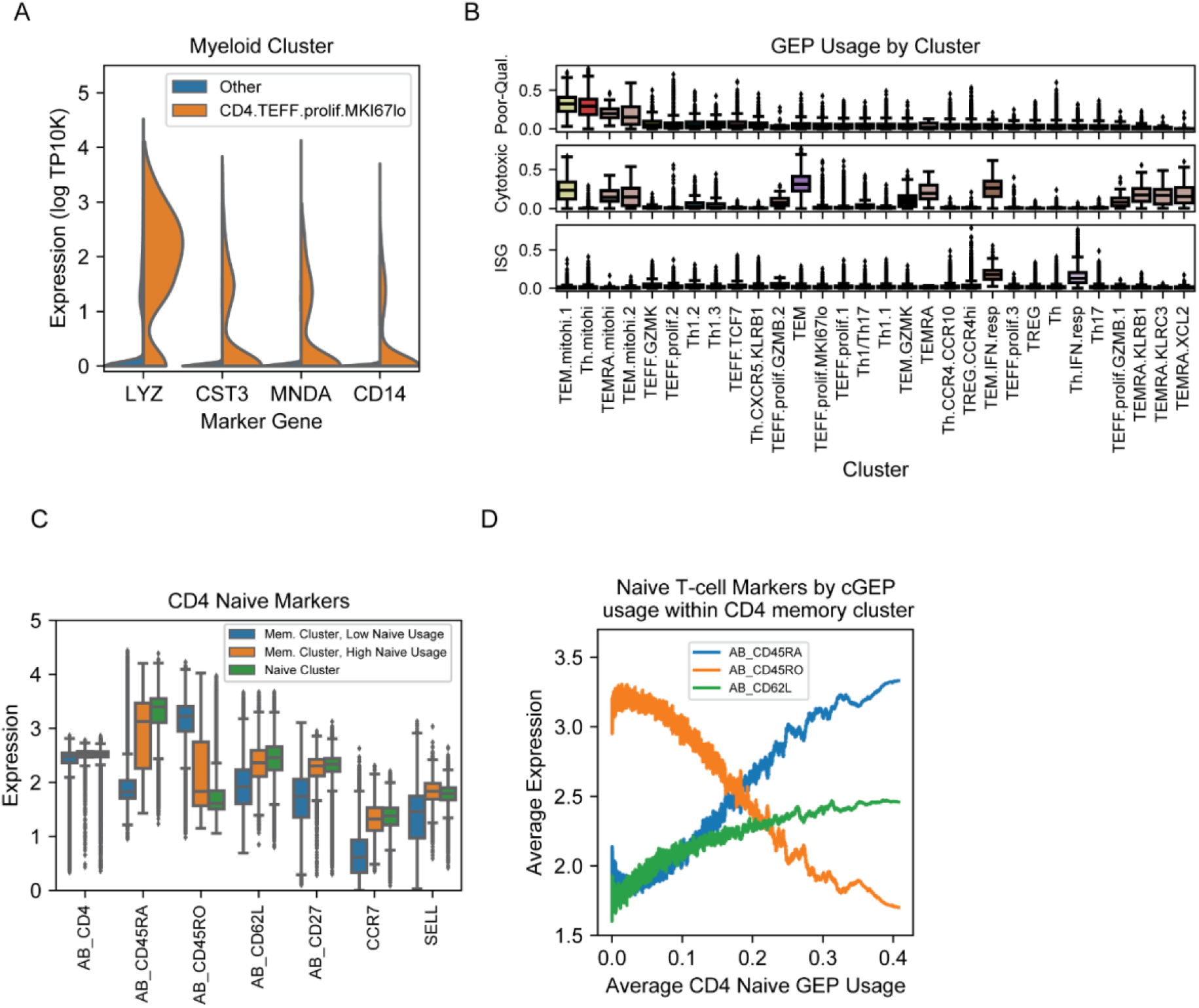
Characterization of COMBAT dataset clustering. (A) Violin plot for myeloid cell marker genes in cells originally annotated as CD4 memory T-cells broken out by the CD4.TEFF.prolif.MKI67lo subcluster, or all other subclusters combined. (B) Usage of the ISG, Cytotoxic, and Poor-quality cGEPs in cells stratified by their CD4 memory subcluster. (C) Expression of CD4 naive marker genes in cells initially clustered as CD4 memories (blue and orange boxes) or CD4 naives (green cluster). Cells initially clustered as CD4 memory are stratified by their usage of the CD4 naive cGEP with a threshold of 0.1.

**Supplementary item 5.**
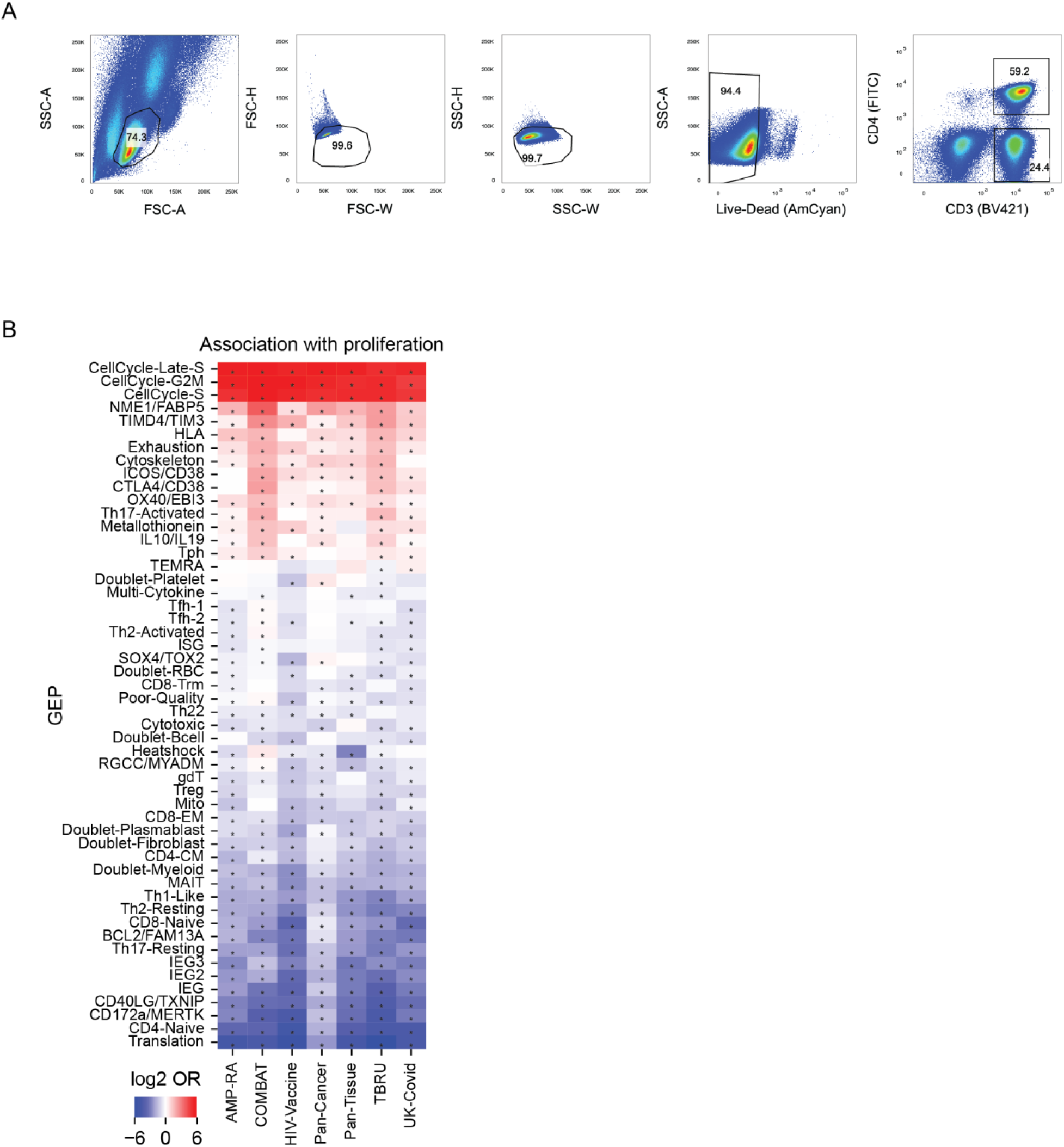
cGEP associations with proliferation across datasets. (A) Gating strategy to identify CD3+ CD4+ and CD3+ CD4-populations in the AIM-Seq experiment. (B) Heatmap of the average Log2 ratio of mean usage in proliferating cells (usage>0.1 of proliferation GEPs) and non-proliferating cells (usage<0.1) for all GEPs (rows) and datasets (columns). An absolute value ceiling of 6 is used to aid visualization. * indicates paired t-test P<.05.

**Supplementary item 6.**
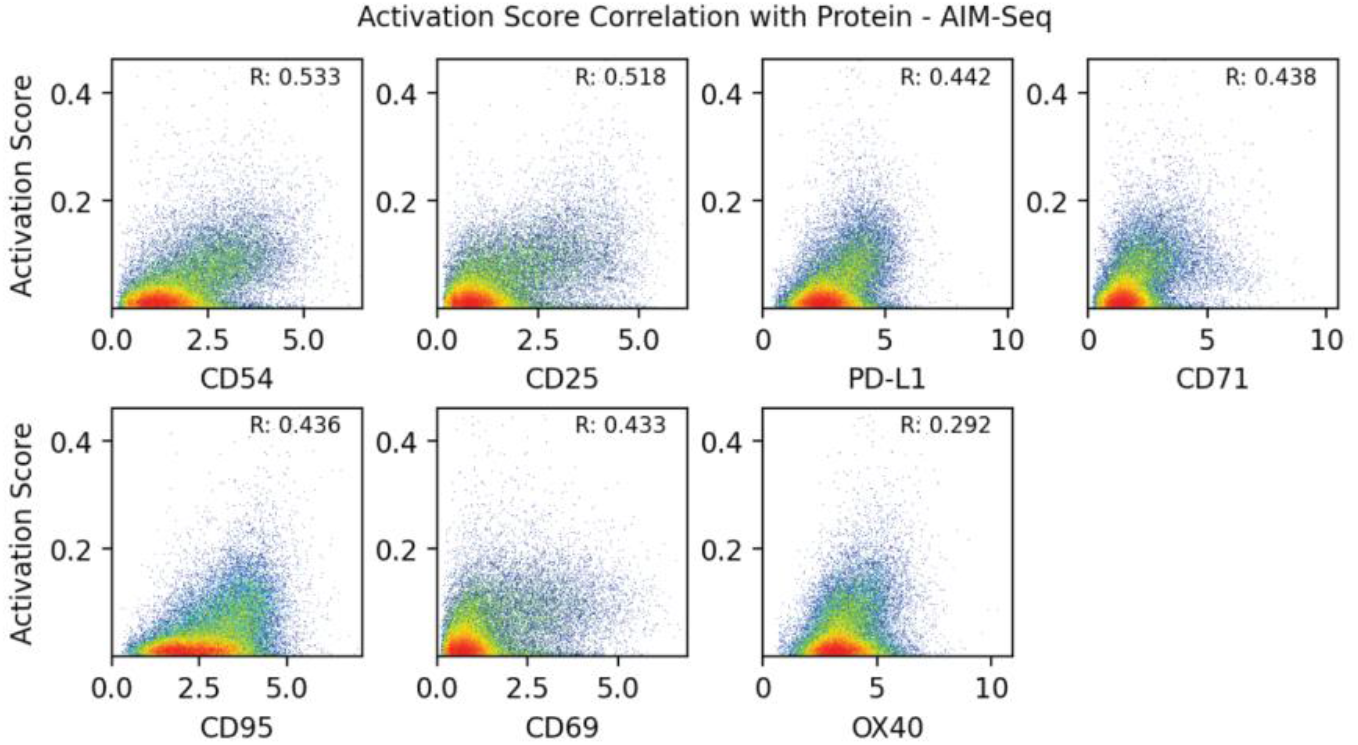
Antigen-specific activation (ASA) score correlation with surface protein activation markers in the AIM-Seq dataset.

